# Dimerization mechanism of an inverted-topology ion channel in membranes

**DOI:** 10.1101/2023.01.27.525942

**Authors:** Melanie Ernst, Esam A. Orabi, Randy B. Stockbridge, José D. Faraldo-Gómez, Janice L. Robertson

## Abstract

Many ion channels are multi-subunit complexes with a polar permeation pathway at the oligomeric interface, but their mechanisms of assembly into functional, thermodynamically stable units within the membrane are largely unknown. Here we characterize the assembly of the inverted-topology, homodimeric fluoride channel Fluc, leveraging a known mutation, N43S, that weakens Na^+^ binding to the dimer interface, thereby unlocking the complex. While single-channel recordings show Na^+^ is required for activation, single-molecule photobleaching and bulk Förster Resonance Energy Transfer experiments in lipid bilayers demonstrate that N43S Fluc monomers and dimers exist in dynamic equilibrium, even without Na^+^. Molecular dynamics simulations indicate this equilibrium is dominated by a differential in the lipid-solvation energetics of monomer and dimer, which stems from hydrophobic exposure of the polar ion pathway in the monomer. These results suggest a model wherein membrane-associated forces induce channel assembly while subsequent factors, in this case Na^+^ binding, result in channel activation.

**Teaser:** Membrane morphology energetics foster inverted-topology Fluc channels to form dimers, which then become active upon Na^+^ binding.

## Introduction

Ion channels enable high-throughput permeation of ions across the cell membrane, typically with exquisite selectivity. To accomplish these seemingly incongruent tasks, a protein structure must feature a stable pathway for the ion offering precise chemical coordination as well as second-shell favorable electrostatic interactions, which together counter the otherwise prohibitive cost of traversing the low-dielectric core of the lipid bilayer. Despite significant architectural variability, the majority of known channel structures show that this typically involves formation of an oligomer where the ion conduction pathway is buried along the central axis; examples include voltage-gated potassium channels, transient receptor potential channels and nicotinic acetylcholine receptors (*1–4*), among many other families.

The assembly of multimeric ion channels into functional units is therefore a ubiquitous process of essential importance in cellular physiology. Yet, the underlying molecular mechanisms remain poorly understood, as protein oligomerization reactions in membranes have been, generally speaking, very challenging to characterize. Fundamental insights have been gained in a few cases, however. Consider the channels formed by gramicidin peptides, which form continuous Na^+^ selective pores upon dimerization of two β-helix subunits, one in each bilayer leaflet (*5*), in a reversible process that is thermodynamically linked to environmental parameters such as the hydrophobic thickness of the surrounding membrane (*6, 7*). Aside from model systems, the only integral membrane transport protein whose assembly thermodynamics has been characterized is the CLC-ec1 Cl^-^/H^+^ antiporter, a homologue to voltage-gated chloride ion channels in humans. This transporter has been shown to exist in a reversible homo-dimerization equilibrium in membranes (*8*), largely controlled by differentials in the lipid solvation energetics of the monomeric and dimeric states (*9*). Reversible oligomerization of membrane proteins occurs also beyond the channel family; a remarkable example is the bacterial flagellar motor, whose membrane-embedded subunits have been shown to be in continuous exchange, even when the motor is active (*10*). Clearly, there is a growing body of evidence showing that many membrane proteins exist in a thermodynamic balance between pseudo-stable monomers and multimers, with both forms adapted to the hydrophobic environment of the membrane, though not identically. Nonetheless, what is particularly perplexing about many oligomeric ion channels is that the polar surfaces lining their central ion pores would be exposed to the hydrophobic core of the lipid bilayer in the monomeric form, seemingly implying only the fully assembled oligomer state is viable in the membrane. Are the mechanisms and controlling factors for the assembly of ion channels fundamentally different from those at play for other membrane protein complexes?

To begin to address this important fundamental question, we developed a new model system for studying ion channel subunit assembly in membranes, namely the inverted-topology, homodimeric ion channel Fluc. The Fluc family consists of small, 4 transmembrane helix membrane proteins that dimerize to form highly selective fluoride (F^-^) channels. These proteins protect unicellular organisms from F^-^ accumulation in the cytoplasm, which inhibits enolase and pyrophosphatase, two enzymes essential for glycolytic metabolism and nucleic acid synthesis (*11, 12*). Flucs exhibit a remarkable >10^4^-fold, F^−^/Cl^−^ selectivity and single-channel recordings show that they are almost always open, *P*_*open,WT*_ > 95% (*13*). Flucs are also the only channels discovered so far to adopt an inverted-topology dimer arrangement. While Flucs are small, the crystal structure of the *Bordetella pertussis* homologue (Fluc-Bpe) shows a large dimerization interface with a surface area per subunit of 1700 Å^2^ (*14, 15*). Along the dimerization interface are two F^-^ permeation pathways with resolvable electronegative densities marking transient binding sites for F^-^ ions (*16*). In addition, there is an unusual electron density well removed from the F^-^ permeation pathways at the dimerization interface and at the very center of the lipid-bilayer core. The tetrahedral coordination formed by the backbones of G77 and T80 from each subunit indicates this density is as a sodium (Na^+^) ion residing along the protein’s two-fold symmetry axis (**Fig. 1A**) (*17*). The unique location of Na^+^ at the dimerization interface in the membrane core suggests that Na^+^ binding to wild-type (WT) Fluc locks the protein in the dimeric state. Indeed, while Na^+^ is not exchangeable with other cations in WT channels at ambient temperature (*17*), the mutation N43S, which weakens Na^+^ binding by disrupting the hydrogen bonding network behind this central site, introduces reversible Na^+^ titration of F^-^ transport, as well as inhibition by Li^+^ (*17*), indicating N43S Fluc is a practical model system to examine the mechanism of channel assembly.

**Figure 1:**
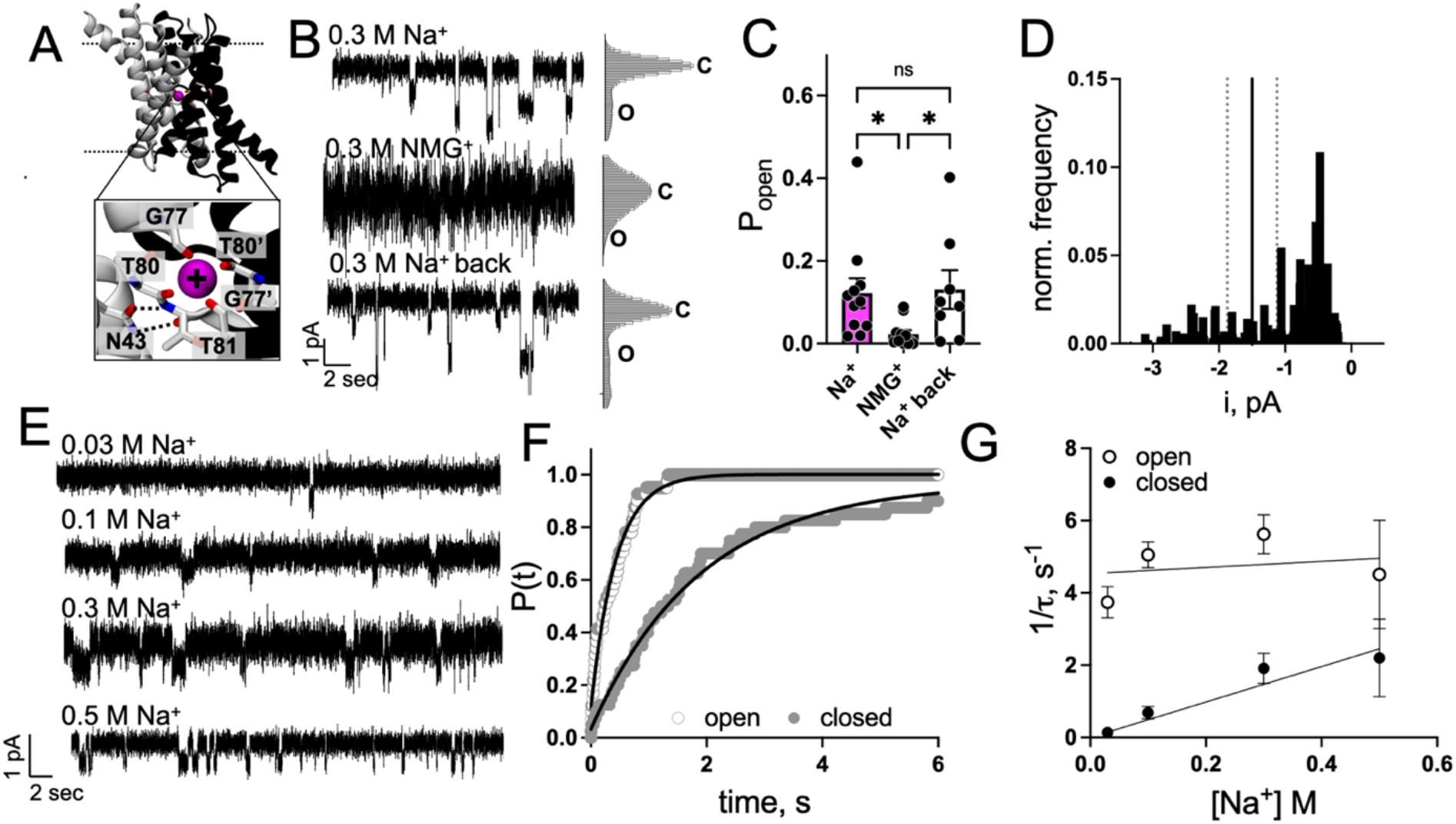
Cyanine5 labeled Fluc N43S-R128C (N43S-Cy5) shows Na^+^ -dependent channel function. (**A**) Na^+^ binding site in Fluc-Bpe (PDB ID: 5NKQ) with residues in coordination distance (G77, T80, G77’, T80’) to Na^+^, N43 side chain which is within hydrogen bonding distance of backbone carbonyl group and backbone amide of T81. (**B**) Na^+^ exchange in single channel recording. Representative trace of single channel recordings of N43S-Cy5 reconstituted at 0.05 μg/mg in EPL, inserted into a 2:1 POPE/POPG bilayer and recorded at -200 mV. Opening of the channel is shown downwards. Single channel was recorded in 300 mM NaF buffer in cis and trans chamber (Na^+^). Then buffer was switched to 300 mM NMGF in cis chamber (NMG). Buffer was then switched back to 300 mM NaF in cis chamber (Na^+^ back). Histograms on the right of the traces show the current amplitude distribution of the trace. (**C**) Probability of channel opening calculated from 8-11 channels from two independent protein purifications (*p-*values: Na^+^ vs NMG^+^: 0.0149, NMG^+^ vs. Na^+^ back: 0.0179; Na^+^ vs. Na^+^ back: 0.8825, unpaired t-test). (**D**) Open state current (i) amplitude histogram of N43S-Cy5 channels recorded at -200 mV holding potential at 300 mM Na^+^. Shown is the normalized frequency of the open states of 28 recorded channels from two independent protein purifications. The solid line marks the WT current amplitude and the dotted line mark a 25% error (*16*). Zero current is defined as the mean level of the fully closed channel. (**E**) Single channel recordings with indicated concentrations of Na^+^ with conditions as in B). (**F**) Cumulative distributions *P(t*) of closed (solid points) and open (open points) dwell times for a single N43S-Cy5 channel in the presence of 0.5 M Na^+^. Solid lines show single-exponential fits. (**G**) Dependence of closed (solid points) and open (open points) time constants on Na^+^ concentration [Na^+^]. Each point represents mean ± SE time constant from three to five separate bilayers. Solid lines show fits to equation (5) and (6) for closed and open intervals, respectively. *k*_*on*_= 4.9 s^-1^ M^-1^ and *k*_*off*_ = 4.5 s^-1^.

Indeed, through a series of experimental and computational investigations, we report that the dimerization of Fluc N43S in membranes follows a thermodynamic model where dissociated subunits exist as pseudo-stable forms in equilibrium with the dimer complex. Single-channel recordings show that Fluc N43S is an ion channel, akin to WT Fluc except that *P*_*open*_ depends on Na^+^. While Na^+^ binding is required for activation, single-molecule photobleaching analysis reveals that the protein dimerizes with strong affinity, *ΔG°* = -10.3 ± 0.4 kcal/mole, 1 subunit/lipid standard state, in the absence of Na^+^; Na^+^ binding to the dimer state is only weakly stabilizing. Coarse-grained molecular dynamics simulations of the Fluc monomer in membranes, shows that the lipid bilayer forms a thinned and curved defect around the dimerization interface, similar to what we observed previously in our studies of CLC (*9*). Finally, simulations of freely diffusing monomers in the membrane, designed to preclude direct protein-protein contacts, nevertheless show that Fluc selectively forms stable dimers in the native orientation, which bury the membrane defect. Taken together, these studies demonstrate that the ion channel N43S Fluc exists in a monomer-dimer equilibrium that appears to be strongly influenced by the morphological energetics of the membrane. We propose that this thermodynamic mechanism is shared by other ion channels, and that it enables the cell to dynamically regulate ion channel assembly, function, and degradation through variations in composition and physical state of the lipid bilayer.

## Results

### N43S Fluc is a Na^+^ dependent ion channel

As mentioned, previous structural studies of WT Fluc revealed a Na^+^ ion bound at the center of the dimerization interface. This ion remains associated with the channel throughout purification, indicating the structure of the dimer interface is highly stable. Only when elevated temperatures are applied to proteoliposomes for extended times can Na^+^ be exchanged with Li^+^, but full removal of Na^+^ results in precipitation (*17*); these observations suggest that passive dissociation of WT dimers in *in vitro* conditions will be rarely observed, and hence that WT Fluc is impractical as a model system. Instead, we focused on N43S Fluc (**Fig. 1A**), a mutant known to weaken Na^+^ binding, reasoning that it would also be more conducive to spontaneous dimer dissociation. However, before proceeding to investigate the dimerization of N43S Fluc, we sought to ascertain that this variant retains the structural and mechanistic features of the WT channel, and is therefore a valid model system. N43S causes F^-^ efflux from proteoliposomes to depend on the Na^+^ concentration (*17*), but intriguingly it also limits the maximal efflux rate to about 2660 ions per second at saturating Na^+^ (*17*). This efflux rate is much smaller than that of WT Fluc (∼10^7^ F^−^ ions/sec at −200 mV (*13*)), and more aligned with those observed for secondary-active transporters, which have more complex conformational mechanisms (typically 100 – 10,000 ions per second, with an estimated theoretical limit of 100,000 (*18, 19*)). This finding could be interpreted as indicative that N43S Fluc is structurally or mechanistically distinct from WT Fluc; however, it is also plausible that the mutation only influences Na^+^ binding and that N43S Fluc functions as a channel, conducting fluoride ion at a rate comparable to WT, but with a lower open-state probability, or *P*_*open*_, explaining the slower macroscopic efflux.

To resolve this question conclusively, we carried out single-channel electrophysiological recordings of N43S Fluc reconstituted in *E. coli* polar lipid (EPL) and fused into 2:1 POPE/POPG bilayers. Indeed, recordings of Cyanine-5 labeled N43S (N43S-Cy5) at saturating Na^+^ (300 mM) showed single-channel activity (**Fig. 1B**), but with different gating behavior. WT Fluc-Bpe exhibits a *P*_*open*_ > 0.95 and a single-channel conductance of 5 pS with few closures or sub-conductance states (*14, 16*). However, at 300 mM Na^+^, N43S-Cy5 Fluc is mostly non-conducting, exhibiting a *P*_*open,Na*_ = 0.12 ± 0.04 (mean ± SE, n = 11) (**Fig. 1C**). Further, we observe variability in channel conductances, with the most observed population exhibiting ∼2.5 pS (∼ -0.5 pA at -200 mV), about half of the WT conductance (**Fig. 1D**). Independent of the conductance, we find the channel gating is Na^+^ dependent in a reversible manner (**Fig. 1BC, Fig. S1**). After exchange of the cis chamber solution from 300 mM NaF to 300 mM n-methyl-d-glucamine (NMG)-F, the channel openings disappear, *P*_*open,NMG*_ = 0.02 ± 0.01 (mean ± SE, n = 11), returning with the addition of 300 mM NaF buffer back to the chamber, *P*_*open,Na-back*_ = 0.13 ± 0.05 (mean ± SE, n = 8). The dependency of *P*_*open*_ on Na^+^ is resolvable in the single-channel recordings (**Fig. 1E**), and dwell-time analysis of open and closed states show that they are exponentially distributed with time constants *τ*_*o*_ and *τ*_*c*_, respectively (**Fig. 1F**). Plotting 1/*τ* as a function of Na^+^ concentration (**Fig. 1G**), we find that the open times are Na^+^ independent, while the closures decrease with increasing Na^+^, supporting a bimolecular binding scheme with Na^+^ binding rate constants of *k*_*on*_ = 4.9 s^-1^ M^-1^ and *k*_*off*_ = 4.5 s^-1^. These experiments demonstrate that N43S-Cy5 Fluc is a Na^+^-dependent ion channel, and that the slow fluoride efflux rates are primarily explained by a much decreased *P*_*open*_, relative to WT, while the single-channel F^-^ conductance is more modestly affected.

### N43S Fluc exists in a dimerization equilibrium in the membrane independent of Na^+^

Having confirmed that N43S Fluc is an ion channel where *P*_*open*_ is Na^+^ dependent, we then investigated whether dimerization and channel opening are concurrent, as is observed for gramicidin channels (*6, 7*). To examine this question, we measured the dimerization reaction of N43S-Cy5 Fluc in EPL bilayers in the absence of Na^+^. Previous studies used dialysis to remove Na^+^ from proteoliposome samples and exchange it with NMG (*17*). While this method is effective, as evidenced by loss of N43S-Cy5 function (**Fig. S2**) we note that traces of Na^+^ may remain as Na^+^ is a major contaminant in many salts, and so we refer to this condition “0” mM Na^+^.

To measure monomer-dimer equilibrium in membranes, we reconstituted N43S-Cy5 in EPL at protein densities ranging from *χ* = 1 × 10^−8^ to *χ* = 2 × 10^−5^ subunit/lipid and measured the photobleaching probability distributions of each sample using the subunit-capture approach (**Fig. 2A**) (*8*). This method recognizes that the photobleaching probability distribution follows a Poisson distribution if one considers heterogenous compartments and multiple protein species and can be modeled using a stochastic simulation approach (*20*). Single-molecule photobleaching analysis requires specific labeling of each subunit and knowledge of the labeling yield. Therefore, we spectrophotometrically measured the site-specific labeling of Fluc N43S at R128C with Cyanine5 via a thiol-maleimide reaction as well as the background labeling of the cysteine-less protein. We found *P*_*site,N43S*_ = 0.67 ± 0.01 (mean ± SE, n = 23), with a non-specific background labeling yield of *P*_*bg*,_ = 0.11 ± 0.03 (mean ± SE, n = 12) (**Fig. S3A**).

**Figure 2.**
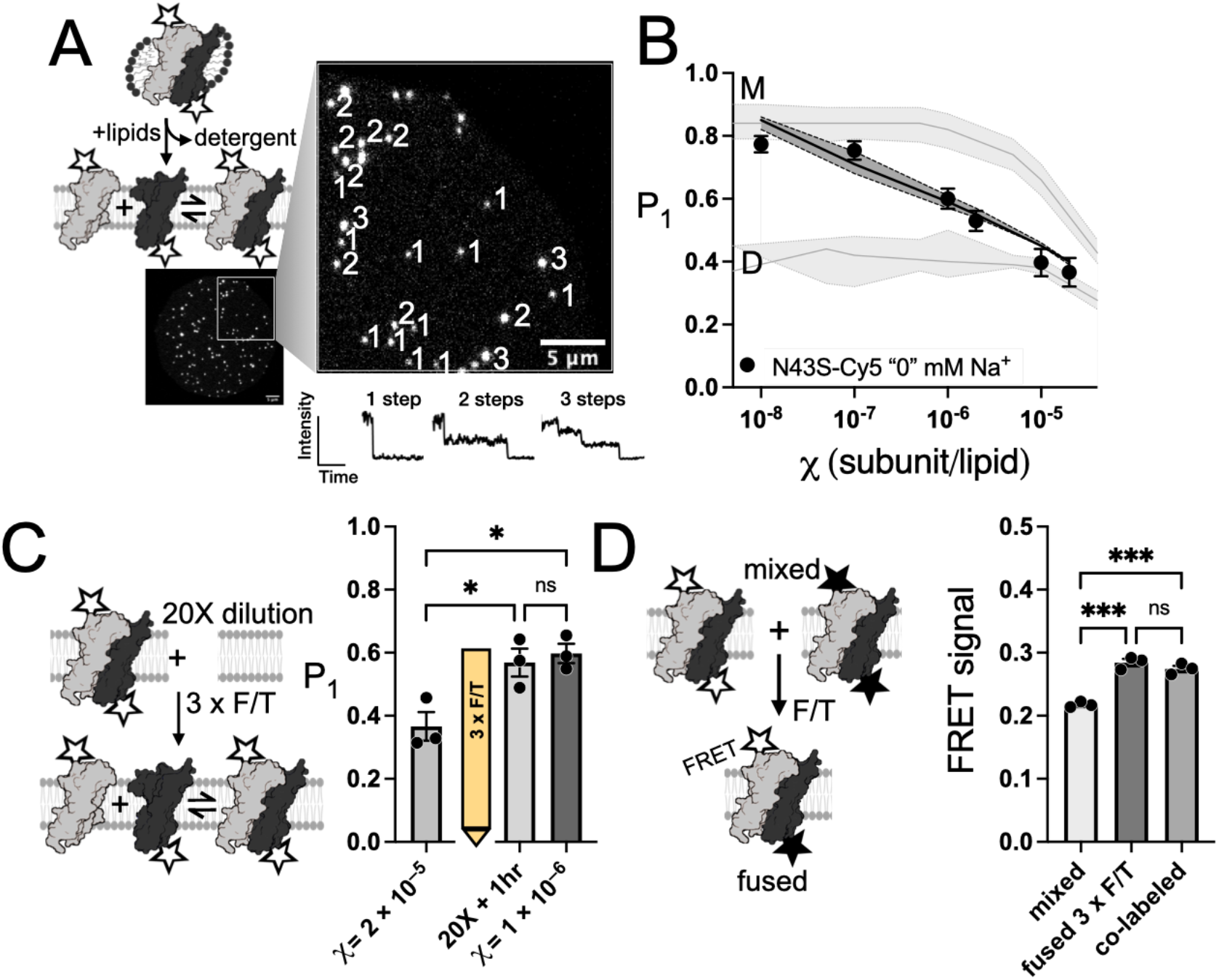
N43S-Cy5 Fluc forms stable dimers without Na^+^. (**A**) Cartoon representation of reconstituting Cyanine5-labeled Fluc N43S (N43S-Cy5) into *E. coli* polar lipid (EPL) bilayers and photobleaching analysis after subunit capture as described by (*8*). Liposomes are imaged under the TIRF microscope and photobleaching steps are counted. **(B**) N43S-Cy5 shows a shift in monomer-dimer distribution over *χ* = 1 × 10^−8^ subunit/lipid to *χ* = 2 × 10^−5^ subunit/lipid. *χ* is the observed mole fraction considering protein recovery yield = 2. Photobleaching probabilities of a single step (*P*_*1*_) vs *χ* for N43S-Cy5 and simulated monomer, *M*, and dimer, *D*, controls. Solid line represents the mean ± SE of the best fit to the experimental data. Data are reported as mean ± SE, n = 3. (**C**) Dilution of reconstituted protein with excess membrane shows a shift in the photobleaching probability distribution. N43S-Cy5 reconstituted at *χ* = 2 × 10^−5^ was diluted by 3 x F/T fusion with 20X excess membrane and incubated at room temperature for one hour (1h). *χ* = 1 × 10^−6^ subunit/lipid shows the *P*_*1*_ for the reconstituted sample. All data are reported as mean ± SE, n = 3 (*p*-values: *χ* = 2 × 10^−5^ vs. 20X + 1hr: 0.0474, *χ* = 2 × 10^−5^ vs. *χ* = 1 × 10^−6^: 0.0270; 20X +1hr vs. *χ* = 1 × 10^−6^: 0.9554, Ordinary one-way ANOVA). (**D**) N43S-Cy3 and N43S-Cy5 undergo rapid subunit exchange. Donor- and acceptor labeled Fluc (N43S-Cy3 and N43S-Cy5, respectively) are reconstituted separately, and liposomes are mixed and fused using repeated F/T cycles (‘mixed’). Upon fusion, subunits can freely exchange leading to an increase in FRET signal over time that approaches a maximal FRET signal determined by colabeling the protein with both fluorophores (‘co-labeled’). Data reported as mean ± SE, n = 3 – 4 (*p*-values: mixed vs. fused 3 x F/T: 0.0001, fused 3 x F/T vs. co-labeled: 0.3425, mixed vs. co-labeled: 0.0003, Ordinary one-way ANOVA).

Comparing the probability of single steps, *P*_*1*_, of N43S-Cy5 to simulated monomer, *M*, and dimer, *D*, species, we observed that the protein population shifts from monomers to dimers as a function of the protein density in the membrane, indicative of an equilibrium dimerization reaction (**Fig. 2B**). If this system truly is in a dynamic equilibrium, then a perturbation by dilution will also yield a shift in the photobleaching probability distribution in the monomeric direction. In lipid bilayers, dilution is not spontaneous, but can be achieved via freeze/thaw fusion of proteoliposome samples with excess membrane. As we “dilute” N43S-Cy5 Fluc at *χ* = 2 × 10^−5^ subunits/lipid with 20 times excess membrane we observe an increase in *P*_*1*_ of the freeze/thawed sample, and this converges with *P*_*1*_ values for a sample that was directly reconstituted at *χ* = 1 × 10^−6^ subunits/lipid (**Fig. 2C**). Finally, as these dilution experiments only report on subunit dissociation, we also assessed the formation of new dimer complexes by monitoring Förster Resonance Energy Transfer (FRET) during subunit-exchange. Here, N43S Fluc dimers are labelled with a donor fluorophore, Cy3, acceptor fluorophore, Cy5, or co-labelled with Cy3/Cy5 and reconstituted into separate EPL membrane samples. At the start of the reaction, the donor and acceptor proteoliposomes are combined into the same membrane by freeze/thaw fusion. Under dynamic equilibrium, dimers will dissociate and then reassemble leading to an increase in FRET signal over time, which is expected to converge with the co-labelled control given comparable Cy3 and Cy5 labeling (**Fig. 2D** and **Fig. S3A-B**). To ensure that the liposomes fully fused, we independently assessed FRET of the membranes themselves using fluorescently labelled lipids, PE-NBD as donor and PE-Rhodamine B as acceptor, doped in preparations of EPL liposomes. The FRET signal saturates after two freeze/thaw cycles and converges to the co-prepared PE-NBD/PE-Rhodamine B positive control (**Fig. S3C**). Therefore, we applied the same approach to the N43S-Cy3 and N43S-Cy5 fusion in “0” mM Na^+^ and observed that the FRET signal increases and converges with the signal from the N43S-Cy3/Cy5 co-labelled positive control after three cycles of freeze/thaw, indicative of new heterodimer formation (**Fig. 2D**). We note that each freeze/thaw cycle takes time, involving 12 minutes of freezing at -80 °C and 15 minutes thawing at room temperature, and so the kinetics of the reaction are difficult to resolve. However, we were able to attain better time resolution by monitoring the time dependency of the FRET signal after a single freeze/thaw cycle, where membranes are about 66% fused. Here, we can observe the exponential increase of the FRET signal that plateaus at a normalized FRET signal of 0.72 ± 0.17 (mean ± SE, n = 3) over the course of the hour with the time constant *τ* = 12.8 ± 3.3 min (**Fig. S3C**). After two additional cycles, the fused sample converges with the Cy3/Cy5 co-labelled control, indicating that N43S Fluc is in a reactive equilibrium with relatively fast kinetics compared to what has previously been reported for CLC dimerization (*8*).

Having verified that N43S Fluc is in a dynamic monomer-dimer equilibrium, we proceeded to quantify the thermodynamic stability of N43S-Cy5 dimer, based on the observed photobleaching probability distributions (**Fig. 2A**). To do this, we used a stochastic simulation of the Poisson process of protein reconstitution into liposomes, with which we modeled the expected photobleaching probability distribution for a reactive monomer-dimer system at a given condition. Given that site-specific and non-specific background fluorophore labeling yields are known (*P*_*fluor*,_ *P*_*bg*_) we can carry out an iterative search of the dimerization dissociation constant *K*_*D*_ that best-fits the experimental photobleaching probability data (*P*_*1*_, *P*_*2*_, *P*_*3+*_). Additionally, we used the cryo-EM liposome size distribution reported by Walden et al. (*21*), and measured the subunit/lipid mole fraction after reconstitution, freeze/thaw and extrusion, which shows that the experimental mole fraction is 200% of that originally reconstituted (**Fig. S4A**). This increase results from a significant loss in lipids (∼ 50%) while the protein recovery is ∼ 100% (**Fig. S4A**). Using the values of *P*_*fluor*_ = 0.67 and *P*_*bg*_ = 0.11 obtained via spectrophotometric analysis, we find the best-fit model yields *K*_*D*_ = 4.0 ± 1.6 × 10^−8^ subunits/lipid (mean ± SE, n = 3), corresponding to *ΔG*° = -10.3 ± 0.4 kcal/mole, 1 subunit/lipid standard state (**Fig. 2A, Fig. S4B**). This demonstrates that N43S Fluc forms dimers even in the absence of Na^+^, with a thermodynamic stability that is comparable to what was reported for CLC-ec1 I422W (*8, 22*).

As a comparison, we also carried out single-molecule photobleaching studies of WT Fluc-Bpe in EPL membranes. For these experiments we switched the labeling site to R29C but found no differences in fluoride transport or labeling yield (**Fig. S5AB**). In contrast to N43S-Cy5, the *P*_*1*_ of ‘WT’-Cy5 overlaps with the dimer model, except for the lowest density condition, *χ* = 2 × 10^−9^ subunits/lipid, which indicates the possible appearance of monomers (**Fig. S5C**). This may correspond to a dimerization reaction that is extremely stable, which our approach cannot measure quantitatively due to the technical limit of diluting the membrane further. Yet, if WT Fluc participates in a reactive equilibrium, then the individual subunits will exhibit dynamic exchange in the membrane over the same time range. To examine this, we fused our ‘WT’-Cy5 proteoliposomes with 3-fold excess of proteoliposomes containing unlabeled protein at the same mole fraction density. If the subunits exchange, then the Cy5 labeling yield will be reduced from 68% to 17%, predicting a shift in *P*_*1*_ = 0.4 to 0.8 upon equilibration. However, we did not observe any change in *P*_*1*_ even after 25 days of incubation at room temperature (**Fig. S5D**), indicating that the WT dimers remained intact over this time. Finally, we also considered whether the histidine tag may influence the dimerization reaction, but the photobleaching probability distributions showed no change after proteolytic his-tag removal (**Fig. S6**). Overall, these results indicate that WT Fluc is a kinetically trapped dimer in the membrane and its thermodynamic stability cannot be measured under our current conditions.

### N43S Fluc dimer stability is only weakly linked to Na^+^ and Li^+^

Having established that N43S Fluc exists in a dimerization equilibrium in membranes in the absence of Na^+^, we then investigated whether Na^+^ is a stabilizing factor for Fluc dimerization using the same single-molecule photobleaching approach presented in **Fig. 2A**, but now titrating the Na^+^ concentration. We found that increasing Na^+^ does promote dimerization, but this is only a small part of the total free energy of dimerization, stabilizing the complex by *ΔΔG* = 1.8 kcal/mole as Na^+^ is raised to 300 mM (**Fig. 3A, Table S1**). These results suggest a model in which Na^+^ binding likely influences the final geometry of the dimer state but is weakly linked to dimerization from a thermodynamic standpoint. A full schematic of possible ligand binding to a dimerizing macromolecule, i.e. Fluc, can be modeled by considering the comprehensive reaction scheme of a monomer-dimer equilibrium in interplay with a ligand, i.e. Na^+^ (*23*) (**Fig. 3B**).

**Figure 3.**
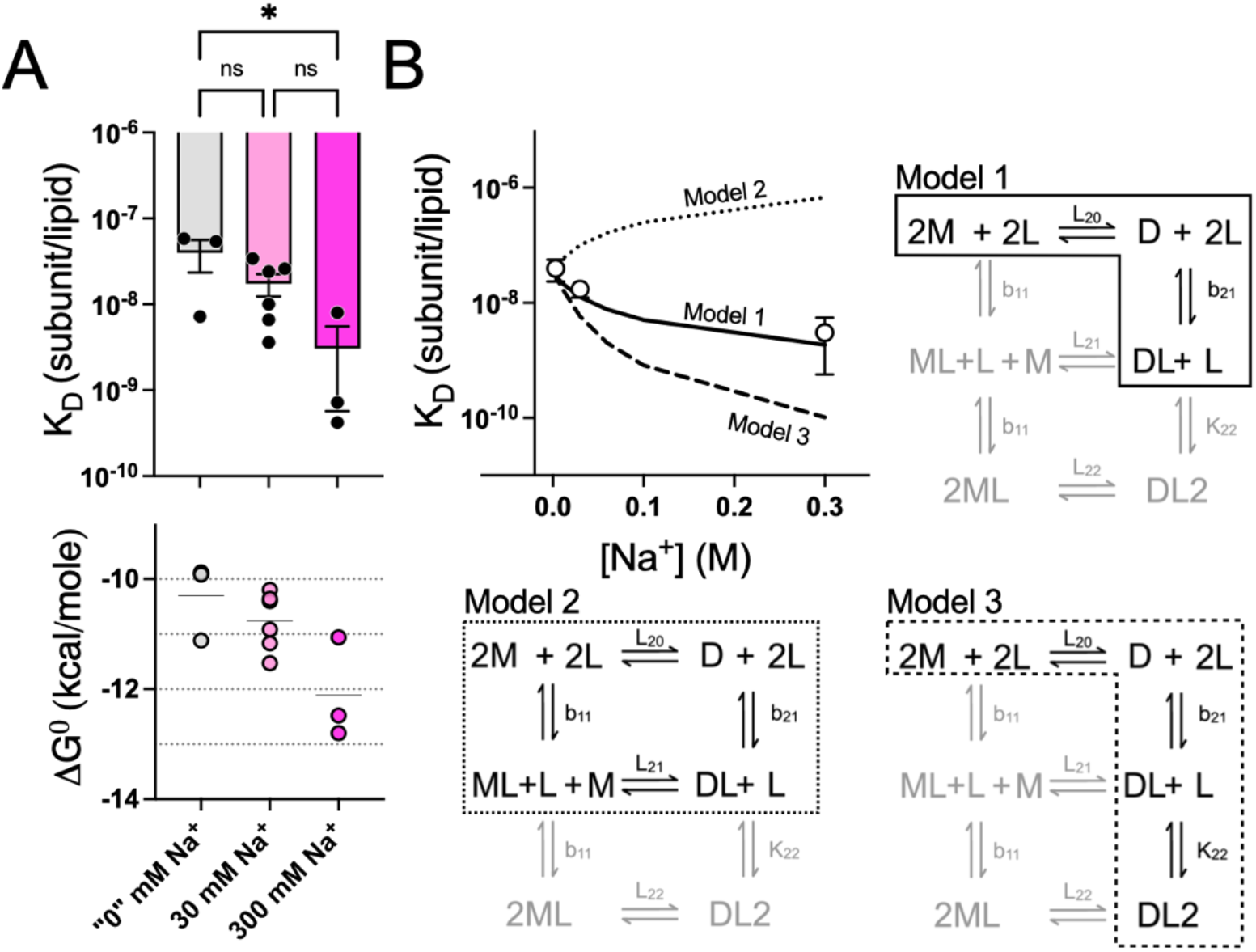
Dimerization provides a single Na^+^ binding site. (**A**) Fluc-N43S dimer is stabilized with increasing concentrations of Na^+^. Fitted *K*_*D*_ following (*35*) and calculated *ΔG°* for “0” mM Na^+^, 30 mM Na^+^, 300 mM Na^+^ (*p*-values: “0” mM Na^+^ vs. 30 mM Na^+^: 0.0840, 30 mM Na^+^ vs. 300 mM Na^+^: 0.2451, “0” mM Na^+^ vs. 300 mM Na^+^: 0.0222, Ordinary one-way ANOVA). (**B**) Na^+^ binding can be described with single binding site to dimer model (Model 1, solid line). Fitted *K*_*D*_ vs. Na^+^ concentration as well as best fit using Model 1 (solid line, *L*_*20*_ = 2.8 ± 0.3 × 10^7^ lipid/subunit, *b*_*21*_ = 60.4 ± 9.6 M^-1^ mean ± std) and simulated data for single binding site to monomer and dimer with equal affinities (Model 2, dotted line) as well as a two independent binding sites to the dimer (Model 3, dashed line). Data were fitted using Scientist^®^ and the models were created using MATLAB^®^. Reaction species array for monomer-dimer reaction with single-site binding of ligand (*L*) to each monomer subunit (*M*) adapted from (*23*). Equilibrium constants beside the arrows apply to those reactions. The reaction between dimer (*D*) and *2L* to form *D*_*2*_*L*_*2*_ is described by the equilibrium constant *b*_*22*_ = *b*_*21*_*K*_*22*_. Sub-reactions of models are indicated with lines. Model 1 (solid line) does not consider *b*_*11*_, or *K*_*22*_; Model 2 (dotted line) does not consider *K*_*22*_; Model 3 (dashed line) does not consider *b*_*11*_.

Modeling the sub-reactions of this system allows us to compare results for different linkage models for the observed Na^+^-dependent stabilization. We first considered the case in which two monomers associate into a dimer and Na^+^ then binds to this species to form the complex captured in experimental structures of WT Fluc (Model 1). We find that this model agrees well with our experimental data (**Fig. 3B**). We estimate the monomer-dimer dissociation constant in the absence of Na^+^ *K*_*D,N43S*_ = *1/L*_*20*_ = 3.6 ± 0.4 × 10^−8^ subunit/lipid and the Na^+^ binding constant *K*_*D,Na+*_ = *1/b*_*21*_ = 16.6 ± 2.7 mM (mean ± std; **Fig. 3, Table S1**). This data agrees well with functional studies obtaining *K*_*D,Na+*_ = 3.6 mM from fitting Fluc-mediated fluoride transport of the mutant to Na^+^ with a single binding site association reaction (*17*). We also considered the alternative case in which a Na^+^ ion binds to each monomer (Model 2). Fitting the data to this model led to no reliable fitting parameters (**Table S1**) so we simulated this reaction assuming that Na^+^ binds with the same affinity to the monomer as we observed for the dimer (*K*_*D,Na+-monomer*_ = *K*_*D,Na+-dimer*_ = 16.6 mM). The simulated model clearly deviates from the experimental data resonating with our unsuccessful fitting efforts (**Fig. 3B**). Finally, while the crystal structure shows a single Na^+^ in the center of the dimer, we wanted to investigate if the observed stabilization effect of Na^+^ is due to specific binding to a single site. For this, we consider a model in which two Na^+^ can bind to the dimer (Model 3). However, these fitting efforts also resulted in an unreliable estimation of parameters (**Table S1**). When we simulated the reaction assuming equal binding affinities for both Na^+^ binding events (*K*_*D,1st Na+*_ = *K*_*D,2nd Na+*_ = 16.6 mM), we find that the simulated model deviates from our experimental data (**Fig. 3B**). Together, our comparison of different hypothetical reactions schemes supports a model where Na^+^ linkage to the dimerization reaction occurs via a single Na^+^ binding to the dimer state.

In addition to Na^+^, Li^+^ has been shown to inhibit fluoride transport in N43S Fluc and is hypothesized to compete with Na^+^ for the central binding site but render it non-functional (*17*) (**Fig. 4A**). We investigated the effect of Li^+^ on N43S Fluc dimerization and observed a concentration dependent stabilization, comparable to our observations with Na^+^ (**Fig. 4B**). The linkage of Li^+^ binding to dimerization can also be described with a single binding site to dimer model similar to our findings for Na^+^ (**Fig. 4C**) and we find *K*_*D,N43S*_ = 3.4 ± 0.2 × 10^−8^ subunit/lipid and *K*_*D,Li+*_ = 13.8 ± 1.6 mM (mean ± std). Furthermore, the results agree with previously reported data from functional studies, *K*_*D,Li+*_ = 0.96 mM when fit to a single binding site association reaction and in the presence of 10 mM Na^+^ (*17*). Thus, this supports a model where Li^+^ also binds to the central site in Fluc, but it does not stabilize a conductive form of the channel.

**Figure 4.**
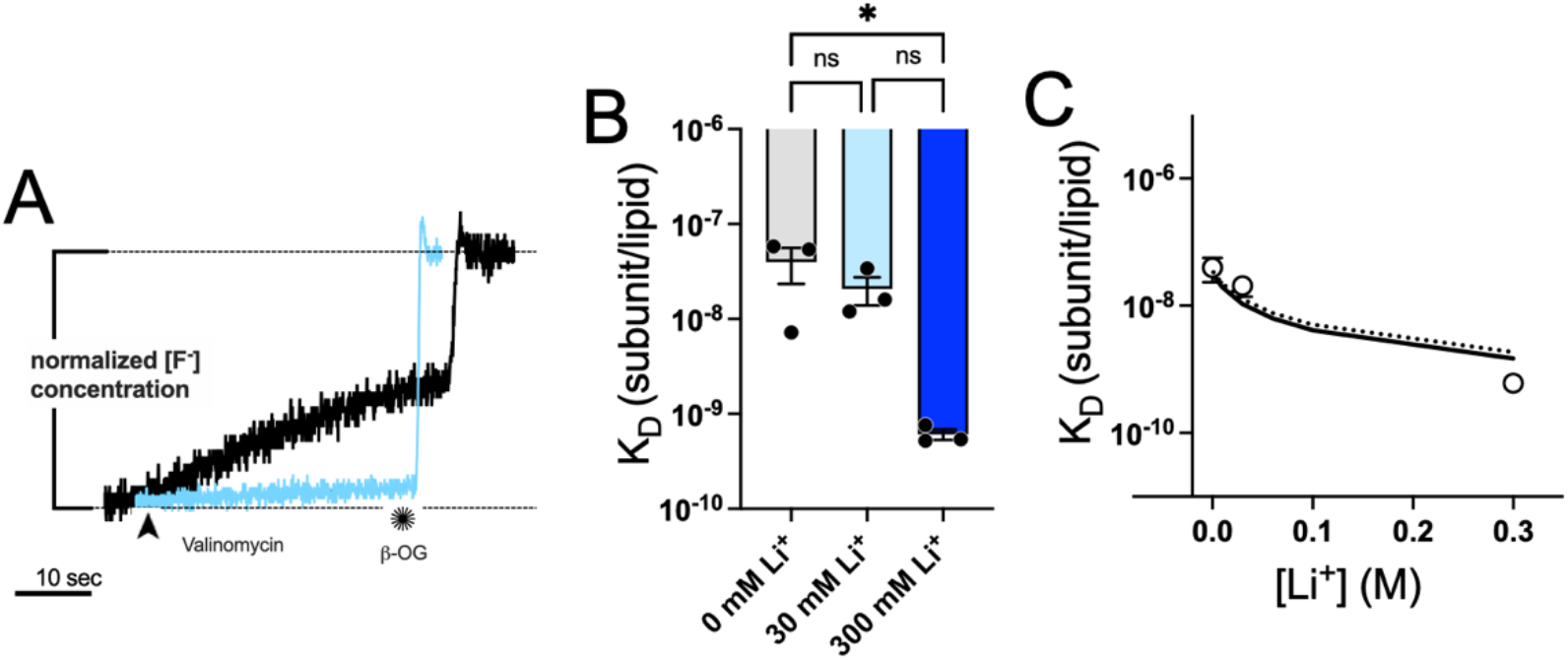
Li^+^ competes for the Na^+^ binding site with similar affinity. (**A**) Competition assay of fluoride export in 10 mM Na^+^ and 30 mM Li^+^ (blue) and with 30 mM Na^+^ added to the sample (black). (**B**) Li^+^ stabilizes dimer state. Fitted *K*_*D*_ following (*35*) for 0 mM Li^+^, 30 mM Li^+^, 300 mM Li^+^ (*p*-values: 0 mM Li^+^ vs. 30 mM Li ^+^: 0.2341, 30 mM Li ^+^ vs. 300 mM Li ^+^: 0.2135, 0 mM Li ^+^ vs. 300 mM Li ^+^: 0.0349, Ordinary one-way ANOVA). (**C**) Li^+^ binding can be described with single binding site to dimer model. Fitted *K*_*D*_ vs Li^+^ concentration aligns well with single site to dimer model (solid line, *L*_*20*_ = 2.97 ± 0.21 × 10^7^ lipid/subunit, *b*_*21*_ = 72.2 ± 8.4 M^-1^, mean ± std) and shows similar affinity as Na^+^ binding (dotted line).

Finally, all stability measurements reported so far were measured in the presence of 300 mM KF, presenting the possibility that the dimerization is dependent on K^+^ or F^-^ in the solution. However, replacing K^+^ with the larger NMG^+^ shows no significant effect, indicating that K^+^ is also too large to fit in the Na^+^ binding site (**Fig. S7**). Furthermore, comparing the dimer stability in 300 mM KCl to 300 mM KF on 30 mM Na^+^ background, we find no significant effect of F^-^ in our system (**Fig. S7**).

### Fluc dimerization is coupled to membrane morphology energetics

Our experiments demonstrate that N43S Fluc forms stable dimers in membranes, and that this occurs before Na^+^ binding, i.e. prior to the formation of the tight complex captured in crystal structures of the active Fluc dimer. This observation implies that there is a driving force that sustains a metastable encounter complex of Fluc monomers in the membrane, sufficiently long-lived to permit Na^+^ binding and thereby channel activation. To investigate this mechanism and the nature of this driving force, we first carried out coarse-grained molecular dynamics (CGMD) simulations of monomeric and dimeric Fluc in 2:1 POPE/POPG membranes. The calculated trajectories are 50-μs long, which allows for complete mixing of the two lipid species as well as multiple turnovers of all lipid molecules in the protein solvation shells in exchange for other lipids in the bulk (*9*). Inspection of the membrane morphology in these trajectories, through 2D maps of the hydrophobic thickness, shows the monomer induces a thinned-membrane defect at the exposed dimerization interface, which disappears upon dimerization (**Fig. 5A**). 3D density maps further reveal that this deformation develops to allow hydration of the exposed charged/polar residues that ultimately form the Na^+^ binding site (**Fig. 5B**). Interestingly, these deformations occur not by way of lipid-chain compression (**Fig. 5C**), but rather through tilting (**Fig. 5D**) and increased entanglement between the two leaflets (**Fig. 5E**). These defects are entirely absent for the dimer. Thus, while the protein surface appears to be suitably solvated for both monomer and dimer, adequate solvation of the monomer requires a significant non-bilayer defect, i.e. an energetic penalty in the dissociated state. We thus reasoned that this membrane-dependent energetic penalty is what stabilizes the encounter complex of Fluc monomers prior to formation of the active dimer form.

**Figure 5.**
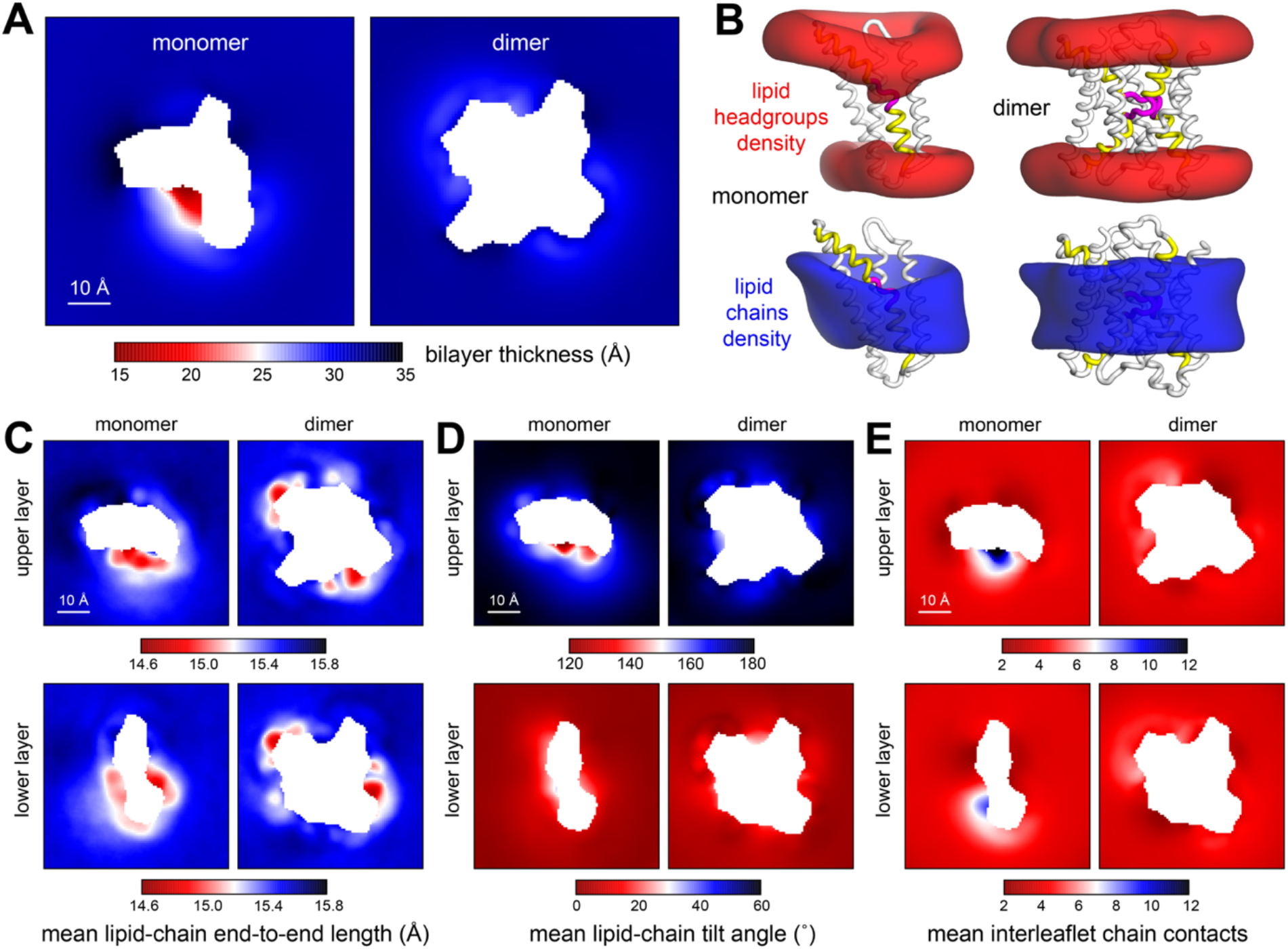
Membrane morphology around Fluc monomer and dimer from molecular dynamics simulations. Results are shown for 50-μs trajectories of either state in POPE/POPG membranes, analyzed using MOSAICS (*39*). All simulations are based on the coarse-grained MARTINI force field. **(A)** 2D map of the local bilayer thickness, quantified by the separation between ester layers in either leaflet. The thinned defect in the monomer marks the location of the dimerization interface. **(B)** 3D density maps for the headgroups (red) or acyl chains (blue) in the vicinity (10 Å) of the protein (white cartoon) for monomer and dimer. The dimerization interface is indicated in yellow, and the Na^+^ binding site in magenta. (**C-E**) 2D maps of the mean lipid-chain end-to-end length, the mean lipid-chain tilt angle (relative to the bilayer perpendicular), and mean number of leaflet-leaflet contacts, for both the upper and lower leaflets of the bilayer. In (C), we detect only marginal changes relative to the bulk membrane, which are also uncorrelated with the oligomerization state; by contrast, in (D) and (E) we detect strong perturbations, only for the monomer state and only at the dimerization interface.

To test this hypothesis, we carried out a second set of simulations (also in 2:1 POPE/POPG bilayers), wherein two Fluc monomers in inverted orientations were positioned at least 5 nm away from each other, in four different starting configurations; each system was then equilibrated and simulated for 30 μs. To specifically examine the mediating role of the membrane, we introduced a short-range repulsive force between the two monomers, which effectively precludes the formation of a contact protein-protein interface. In other words, while the monomers are free to diffuse and transiently collide, they are not permitted to form direct protein-protein interactions in any orientation (an approach that also enabled us to avoid known deficiencies in the CG representation of this kind of interactions (*24*). Over the course of the trajectories, we observed multiple reversible collisions between monomers in non-native orientations. However, when the two monomers happened to approach each other oriented as in the native dimer, i.e. in a manner that aligns the corresponding membrane defects, the monomers formed a non-contact dimer that was stable for the remainder of the simulation (**Fig. 6A**). This behavior was reproducible and observed in all replicate simulations. Time-averaged analysis of the lipid configurations in the monomer, non-contact dimer and dimer states reveals the changes in lipid solvation accompanied with association (**Fig. 6B**). While the monomer carries a non-bilayer defect, exhibiting tilted and entangled lipids, the non-contact dimer allows the headgroups to fan out into a toroidal geometry that appears to match the polar/charged features of the Fluc dimerization interfaces (**Fig. 6C**), and is therefore advantageous. Ultimately, full dimerization requires expulsion of all these lipids, where they can return to their preferred bilayer state (**Fig. 6D**). Altogether, these results demonstrate that differences in membrane morphology and solvation energetics can physically drive Fluc monomers into a non-contact dimer with the native orientation, prior to Na^+^ binding and the formation of a tight protein-protein interface suitable for F^-^ permeation.

**Figure 6.**
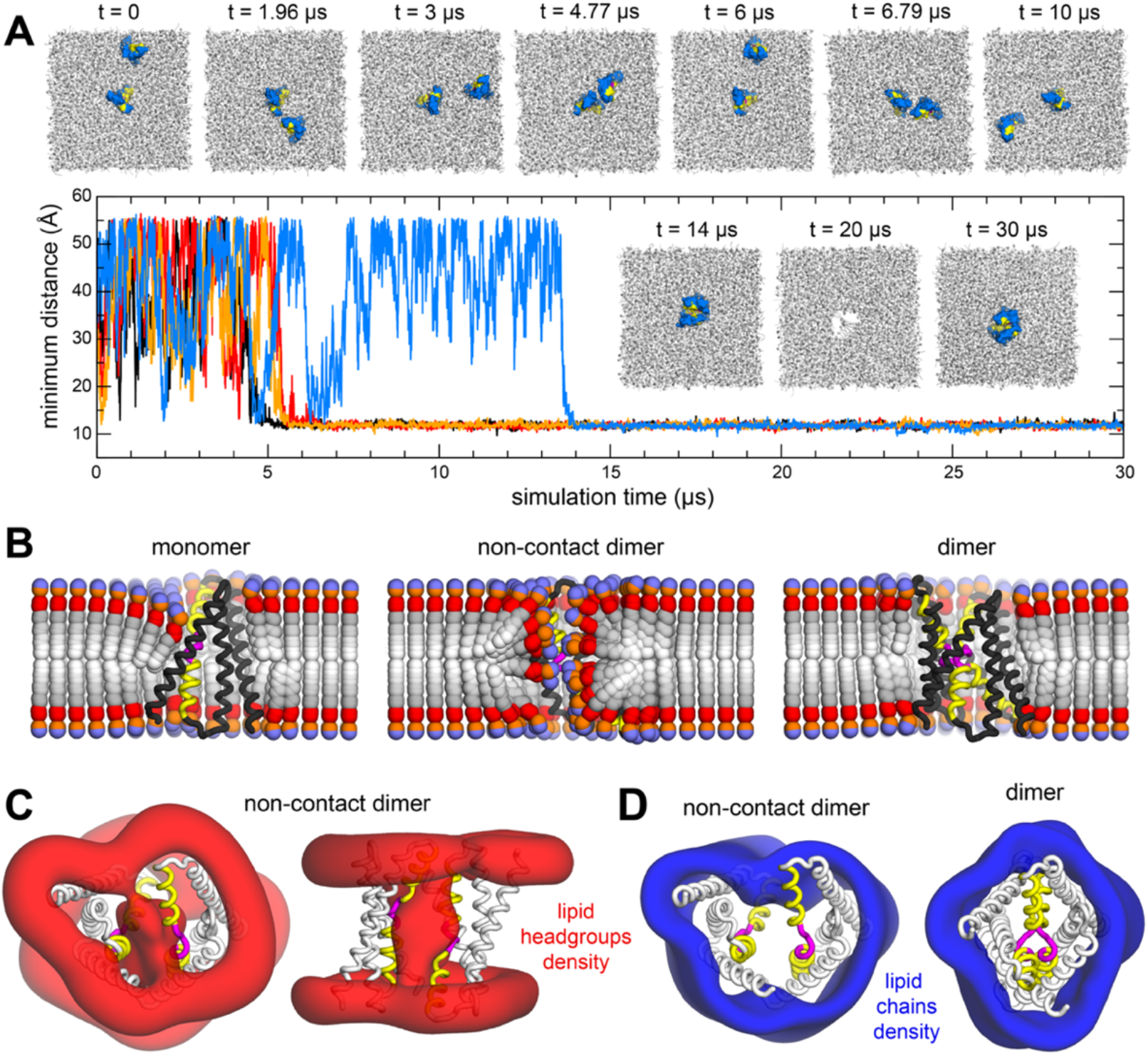
Fluc dimerization is coupled to membrane morphology energetics. Results are shown for four independent trajectories of 30 μs in POPE/POPG membranes, analyzed using MOSAICS (*39*). All simulations are based on the coarse-grained MARTINI force field. (**A**) When freely diffusing Fluc monomers (marine blue) encounter each other in an orientation that aligns their dimerization interface (yellow) and Na^+^ binding motifs (magenta), they form a lipid-mediated encounter complex, stable even in the absence of protein-protein contacts (which are excluded by design). Other orientations are unstable. Snapshots are shown of one of the trajectories at various time points (*t*), with the protein hidden in the 20 μs snapshot to highlight the layer of lipids that remain between the two subunits. Shown is also the minimum distance between monomers (measured at their backbones) versus simulation time, for each of the 4 independent trajectories, showing the same non-contact complex is ultimately observed in all cases. (**B**) Time-averages of the instantaneous 3D conformation of lipid molecules residing in different positions across the membrane plane. Acyl chains (gray scale), ester linkages (red), and headgroups (orange/purple) are shown as spheres. Note perfectly isotropic dynamics, when time-averaged, results in a linear structure for the entire molecule, perpendicular to the membrane mid-plane, and with both acyl chains superposed. These structures are therefore non-physical, but they reveal the mean tilt of the lipid molecules across the membrane as well as the degree of contacts between leaflets. Data is shown for the monomer, for the non-contact dimer (from simulations described in (A)), and for the dimer. (**C**) 3D density maps for the lipid headgroups (red) in the vicinity (10 Å) of the protein (white cartoon) for the non-contact dimer. (**D**) 3D density maps for the acyl chains (blue) in the vicinity (10 Å) of the protein (white cartoon) for the non-contact dimer and for the fully assembled dimer.

## Discussion

### α-helical ion channel subunits can exist in a balance of dissociated and associated states supporting a thermodynamic model of assembly

Our investigation demonstrates that an obligate homomeric ion channel can exist in a monomer-dimer equilibrium in membranes. For Fluc, this becomes apparent once we destabilize the central Na^+^ binding site via the N43S mutation. While this mutation reduces the macroscopic fluoride permeation rate to a value reminiscent of a secondary-active transporter, our electrophysiological measurements demonstrate that N43S Fluc is a *de facto* channel whose single-channel conductance is within 2-fold of the WT protein. We therefore conclude that WT and N43S dimers are structurally and mechanistically comparable, when Na^+^ is bound. For N43S, however, the probability of channel opening is strictly coupled to the Na^+^ concentration. This indicates that N43S Fluc dimers must be conformationally flexible to allow for equilibrium titration of Na^+^, since the ion binds to a site at the core of the dimerization interface (*14*). Indeed, our results for N43S Fluc demonstrate that the removal of Na^+^ does not necessarily involve full dissociation of the monomers, as our single-molecule photobleaching studies demonstrate that the protein dimerizes, and in fact participates in a dynamic monomer-dimer equilibrium reaction with a *K*_*D*_ = 4.0 ± 1.6 × 10^−8^ subunits/lipid, corresponding to *ΔG*° = -10.3 ± 0.4 kcal/mole, 1 subunit/lipid standard state, even when Na^+^ is absent. Therefore, the stability of N43S dimers without Na^+^ is still quite strong. To relate it to a biological reference, this means that if just two N43S Fluc subunits are expressed in an *E. coli* cell, assuming an inner membrane surface area of 4 μm^2^ containing ∼10^7^ lipids, then these subunits would be observed as dimers 70% of the time. It is reasonable to expect that more than two protein copies would be expressed upon activation of the fluoride riboswitch (*25*) and so we predict that even a construct such as N43S would be dimeric in the cellular environment. At 300 mM Na^+^, close to the biological salt conditions, the dimerization would be further stabilized, but only by 1.8 kcal/mole. Together, these factors present the possibility that Fluc monomers may be independently expressed in the membrane, released, and then form stable dimer structures by thermodynamic mechanisms. While it is still open for question if the dynamic assembly is generalizable for other channels and transporters, our studies indicate that the folding of these ion channels may comprise the distinct steps of an oligomerization equilibrium reaction.

### Fluc monomer association eliminates costly lipid-bilayer defects, an emerging theme for membrane protein oligomerization

The observation that N43S Fluc exists in a monomer-dimer equilibrium in membranes, whereby stable but conformationally flexible dimer complexes are formed in the absence of Na^+^, raises the question of what thermodynamic forces sustain this association. To understand this process, we first considered the dissociated Fluc monomer in the membrane. In this state, 1700 Å^2^ of previously buried surface becomes exposed to the hydrophobic core of the membrane for each monomer (*14*), including polar and charged motifs that would otherwise interact with F^-^ or Na^+^. Since exposure of these motifs to the low dielectric of the membrane core is electrostatically unfavorable, the surrounding membrane changes its shape to minimize this hydrophobic mismatch. Our CGMD simulations show that the membrane morphology becomes non-bilayer like, with lipids significantly tilted to facilitate access to hydration. While polar and charged motifs on the protein are now more adequately solvated, this state incurs an energetic strain on the lipid bilayer. From recent studies we know that the morphological energetics of the membrane are non-negligible and must therefore be incorporated into the conceptual models and theories used to describe membrane-protein mechanisms (*26*). Indeed, our simulations of free Fluc subunits, that explore all configurations where protein contacts cannot form, demonstrates that dimerization is overwhelmingly favorable, but only via the native interface. This result demonstrates that it is the membrane energetics, and the relief of the thinned, non-bilayer defect, that drives Fluc to ultimately dimerize in this configuration. This is similar to our findings for CLC dimerization, which binds via a hydrophobic interface that also introduces a thinned, non-bilayer defect in the monomeric state (*9*). Thus, the observation that both Fluc, via a polar/charged interface and CLC, via a greasy interface, have affinity due to their differential membrane structures, presents the membrane as a generalizable driving force for specific, oriented membrane protein assembly in membranes. Note, in Fluc, this stable state does not correspond to the presumed final, functional state, which we assume corresponds to the crystal structures (*14, 16*). This non-contact dimer or encounter complex is mediated by lipid head-groups but is more energetically stable in the membrane than the dissociated monomers and their thinned defect. Therefore, we propose that it is a lipid-solvated stable intermediate along the dimer pathway stabilized by the membrane. Of course, the protein plays a role also, but it is to adopt a stable monomeric structure that imposes a change in the surrounding membrane structure as well as providing precise chemical interactions to form the fluoride selective pore. We note, these types of lipid-solvated non-contact complexes have been observed previously, notably with the ATP synthase, that dimerizes at a distance to stabilize a bent structure of the membrane, promoting self-assembly in rows along the cristae of mitochondria (*27*).

### Na^+^ binding stabilizes the active state and may kinetically trap the dimer

The observation of a non-contact lipid-solvated state allows us to speculate about the mechanism of Na^+^ binding to Fluc channels. Previously, it had been demonstrated that WT Fluc possesses a buried Na^+^ that does not dissociate, unless the protein is heated for an extended time (*17*). Naturally, this raises the question of how Na^+^ accesses its binding site in the first place. One possibility is that the N43S mutation promotes a local conformational change that provides access to the Na^+^ binding site. However, we consider such structural rearrangements unlikely as the channel is formed by two, identical but inverted subunits that would be capable of making the same rearrangements on both sides, thus making it permeable to Na^+^, which is not observed experimentally (*17*). Therefore, based on our findings, we propose that when WT Fluc subunits are expressed in the cellular membrane, they fold into metastable monomeric structures that ultimately associate into dimers, to eliminate the unfavorable thermodynamic cost of lipid solvation of the dissociated state. Dimerization proceeds via the formation of a non-contact lipid-solvated stable intermediate where water and Na^+^ can penetrate the head-group region between the two subunits, but without a continuous pathway across the membrane. With this, the site, formed by the protein backbone, becomes stabilized and assembles around the Na^+^ ion, expelling lipids in the last step to allow for the two subunits to come together in the state that permits highly selective F^-^ permeation. Our studies of the equilibrium linkage of Na^+^, or even Li^+^ binding to dimerization, supports the model that a binding site is present only in the dimer state. The linkage of Na^+^ and Li^+^ binding to dimer stability is weak, and therefore indicates that the ion binding is not a major driving force for dimerization. However, Na^+^ is absolutely required for channel activation, meaning that Na^+^ stabilizes the conductive structure upon binding. Interestingly, Li^+^ presents the same weakly stabilizing effect on dimerization, but inhibits channel function, indicating that it binds but does not stabilize the proper conformation of the channel that allows for F^-^ permeation.

Finally, Na^+^ may have an additional role in WT Fluc, which is to provide a mechanism of kinetic trapping of the active dimer state. We cannot yet measure the thermodynamic affinity of WT dimers, as subunit exchange has not been observed in our hands. However, one way to achieve kinetic trapping is to couple the dimerization to extremely strong affinity of a cofactor. In N43S, the Na^+^ binding affinity is weak (*K*_*D,Na+*_ ∼ mM) but a more favorable binding site (*K*_*D,Na+*_ ∼ nM-μM) would further stabilize the dimerization reaction and make the observation of dissociation improbable. Since the physiological role of Fluc is to allow an immediate leak pathway for F^-^ to leave the cytoplasmic compartment, there must be a significant evolutionary drive to evolve a site that ensures that the channel is always on, but that it also remains “locked” in the assembled state in the membrane. When examining other ion channel structures, it becomes apparent that many channels adopt functional structures that appear to lock an oligomeric assembly in place. Examples include coiled-coiled domains, as in the case of the KcsA K^+^ channel (*28*); cytosolic dimerization domains, as in the CBS domains of eukaryotic CLC channels and transporters; domain-swapping in the membrane, as observed in some of the vanilloid subtype of transient receptor potential channels such as TRPV1 (*29*) and TRPV2 (*30, 31*); or transitioning from an oligomer to a multi-domain structure through evolution as is speculated for Na^+^ and Ca^2+^ channels (*32*). All these examples suggest that there is an evolutionary pressure to keep ion channel complexes kinetically together, which indirectly implies that the membrane embedded subunits have an ability to dissociate within the membrane.

In summary, we discovered that the dimeric fluoride channel Fluc assembles via a dynamic equilibrium reaction of pseudo-stable monomers, driven primarily by membrane-dependent forces. Because the monomers introduce a membrane defect in the dissociated state, monomers that find each other in the correct orientation in the membrane can dimerize, prior to the formation of a protein-protein interface. Na^+^ binding to this encounter complex then brings the protein into its functional state. The stabilization of the functional, conductive channel is dependent on Na^+^ binding, that only offers weak stabilization of the dimer complex. For WT in which Na^+^ binding affinity is presumed to be stronger, the dimer is kinetically trapped in the conductive state. But simply weakening the backbone interactions around this site with N43S, reveals the underlying dynamic and thermodynamically driven reaction. These results map out the fundamentals of ion channel assembly, demonstrating that dynamic complexes can form and can be locked into conformational states with further ion and potential protein interactions. This study thus sets up a general hypothesis for how ion channels assemble and disassemble in biological membranes.

## Methods

### Fluc constructs

Fluc-Bpe bearing two functionally neutral mutations, R29K and E94S, was introduced into a pASK vector encoding a C-terminal LysC recognition site and hexahistidine tag (TRKAASLVPRGSGGHHHHHH). All mutations were made by Quickchange II Site-Directed Mutagenesis (Agilent, Santa Clara #200523) followed by DNA sequencing of the full gene. We introduced two labeling sites, R29C, R128C. The N43S mutation was on the background of the R29K and E94S mutation. The molecular weights for each construct are 15404 g/mol and they have an extinction coefficient *ε*_*Fluc*_ = 39380 M^-1^ cm^-1^. Molecular weights and extinction coefficient were calculated using the Peptide Property Calculator (http://biotools.nubic.northwestern.edu/proteincalc.html).

### Protein Purification

Expression and purification of Fluc-Bpe was carried out as previously described. C41 E. coli competent cells (Lucigen, #60442-1) were transformed with the plasmid and then 2 L Terrific Broth supplemented with ampicillin was inoculated and grown at 37°C. Protein expression was induced with IPTG at OD600 = 1.0. After 1.5 hr of induction, cells were harvested, then lysed by sonication in buffer supplemented with 5 mM reducing agent TCEP (Tris(2-carboxyethyl)phosphine) (Soltec Ventures, #M115) and pH adjusted to 7.5. Protein extraction was carried out with 2% n-Decyl-β-D-Maltopyranoside (DM; Anatrace #D322S25GM) for 2 hr at room temperature. Cell debris was pelleted down and the supernatant was run on a 2 mL column volume (CV) TALON cobalt affinity resin (Takara Bio #635504) equilibrated in CoWB/TCEP (100 mM NaCl (Sigma-Aldrich 746398), 20 mM Tris (RPI #1005479), 1 mM TCEP, pH 7.5 with NaOH, 5 mM DM). After binding, the column was washed with 15 CVs of CoWB/TCEP followed by a low imidazole wash of CoWB/TCEP containing 40 mM imidazole (RPI #I52000). Fluc was eluted with CoWB/TCEP containing 400 mM imidazole, then concentrated in a 10 kDa NMWL centrifugal filters (Amicon #UFC9003) to ∼500 μL and injected on a Superdex 200 10/30 GL size exclusion column (GE Healthcare #28990944) equilibrated in size exclusion buffer (SEB, 150 mM NaCl, 10 mM HEPES (Fisher Scientific # BP310-1), 10 mM NaF (Sigma-Aldrich #201154) pH 7, 5 mM analytical-grade DM (Anatrace #D32225GM)), attached to a medium pressure chromatography system (NGC, Bio-Rad). Protein fractions were collected, and the protein concentration was calculated from the absorbance at 280 nm (A_280_) using a Nanodrop 2000c UV–VIS spectrophotometer (Thermo Scientific).

### Labeling of purified protein

Cyanine3 (Cy3)-maleimide and Cyanine5 (Cy5)-maleimide dye was obtained as lyophilized powder as 50 mg (Lumiprobe #21080 #23080), stored as 10 mM master stocks (50 μl each) in anhydrous Dimethyl sulfoxide (DMSO, Sigma-Aldrich #276855) at -80 °C. The fluorophore conjugation reaction was carried out in SEB with 10 μM Fluc subunits and 50 μM Cyanine3-maleimide, Cyanine5-maleimide, or a 1:8 mixture of both for 15 min at room temperature in dark. At the end of the reaction, 100-fold molar excess of cysteine (RPI #1004738) was added to quench the maleimide reaction, from freshly prepared 100 mM stock in CoWB, pH adjusted to 7.5. The ‘free’ dye was separated from the labeled protein by binding the reaction mixture to a 250 μL cobalt affinity resin column equilibrated with 15 CV CoWB (no TCEP) in a Micro-Bio spin chromatography column (Bio-Rad Laboratories # 7326204), washed 15 CV with CoWB and then eluted with 400 mM imidazole in CoWB, manually collecting only the fluorescently labeled protein. To remove the interfering absorbance of imidazole at 280 nm, the labeled protein was added to a 3 mL Sephadex G50 size exclusion column (Cytiva #GE17-0042-01) equilibrated in CoWB (no TCEP). The fluorescently labeled protein was eluted after addition 2–2.5 mL of CoWB to the column. The concentration and labeling efficiency of protein calculated from the UV-VIS absorbance spectrum of the sample and λ_max_ of Fluc (280 nm), Cy3 (552 nm), and Cy5 (655 nm) as follows:

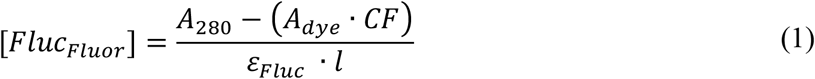

where *l* is the pathlength of 1 cm, *ε*_*Fluc*_ is the extinction coefficient, 39,380 M^-1^ cm^-1^, *A*_*dye*_ is the peak absorbance of the dye (*A*_*Cy3*_ = 552 nm, *A*_*Cy5*_ = 655 nm) and *CF* is the correction factor of the dye absorbance at 280 nm (*CF*_*Cy3*_ = 0.08, *CF*_*Cy5*_ = 0.05).

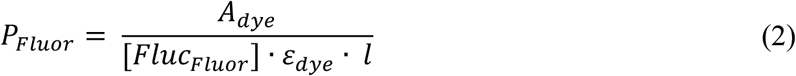

Where *ε*_*dye*_ is the extinction coefficient of the dye (*ε*_*Cy3*_ = 1.5 × 10^5^ M^-1^ cm^-1^, *ε*_*Cy5*_ = 2.5 × 10^5^ M^-1^ cm^-1^). Note, for co-labeled sample, the concentration is calculated using the sum of *A*_*Cy3*_ × *CF*_*Cy3*_ and *A*_*Cy5*_ × *CF*_*Cy5*_. The acceptor to donor labeling ratio is calculated as *P*_*Cy5*_*/P*_*Cy3*_.

### Lipid preparation and protein reconstitution

Reconstitution of Fluc was carried out as described previously (*13*). Briefly, 25 mg/mL chloroform stocks of *E. coli* polar lipids (EPL, Avanti Polar Lipids Inc. #100600C) were dried by evaporation under a stream of nitrogen gas until a thin dried film of lipids appeared. The film was washed with pentane (Sigma-Aldrich #236705-1L). Then, lipids were resolubilized in the required dialysis buffer with 35 mM CHAPS (Anatrace #C316S) for a final concentration of 20 mg/mL lipids. The lipid-detergent mixture was solubilized using a cup-horn sonicator (Qsonica, Newtown, CT) until the sample was transparent. Protein was added to the solubilized lipid-detergent mixture and placed into 3,500 MWCO dialysis cassettes (Thermo Scientific #66330), and then the samples were then dialyzed in the dark at room temperature in 1000x sample volume with 4 buffer changes every 4–12 hours. At the end of dialysis, the samples were freeze-thawed to form large multi-lamellar vesicles (MLVs). This involved 3 repetitions of freezing the samples at -80 °C and thawed in a room temperature water bath. Samples were stored at room temperature, in the dark for the desired amount of time or frozen at -80 °C until used.

### His-Tag Cutting

His-Tag was cut when indicated after reconstitution of labeled protein into EPL lipids. Proteoliposomes were passed through a 0.4 μm polycarbonate membrane (Whatman) using a LiposoFast-Basic extruder before lysine endoproteinase C (LysC, Pierce #90051, 1:20 w/w) was added and sample was incubated for 1 hour at RT. Since Fluc is a dual-topology protein and LysC cannot permeate through the membrane, proteoliposomes were freeze-thawed to form large multi-lamellar vesicles and allow for reorganization of the proteins. This involved 3 repetitions of freezing the samples at -80 °C and thawed in a room temperature water bath. Samples were then again passed through a 0.4 μm polycarbonate membrane using a LiposoFast-Basic extruder before LysC (1:20 w/w) was again added and incubated for 1 hour at room temperature.

### Fluoride transport assay

The fluoride flux assay was performed as described previously (*13, 33*). Here, liposomes containing 300 mM fluoride are placed in a low fluoride solution. Fluoride efflux is initiated by addition of valinomycin (Sigma-Aldrich #V0627) and monitored using a fluoride-selective electrode. After the fluoride efflux reaches a steady state, liposomes are disrupted by the addition of n-octyl-beta-D-glucoside (β-OG, Anatrace #O31125GM) to release the remaining encapsulated fluoride. The change in fluoride concentration over time normalized to the total fluoride concentration in the sample *Δf*_*T*_ can be fitted using Equation 3 adapted from (*21*).

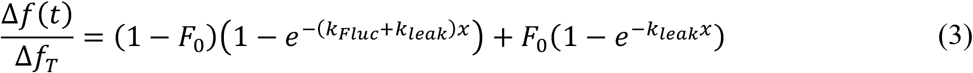

In which *F*_*0*_ is the fraction of liposomes that does not contain active protein and *k*_*Fluc*_ and *k*_*leak*_ are the rate constants for F^-^ flux through the channel and for the background leak through the liposome membrane, respectively. For WT Fluc however, transport rates exceed the response time of the electrode. Therefore, the volume fractions of liposomes with no functional protein (F_0, vol_) are calculated by subtracting the plateau value after addition of β-OG from the plateau value after addition of valinomycin.

### Planar lipid bilayer recordings

Single-channel recordings were performed as described previously via a Nanion Orbit-mini planar bilayer system, using 70% 1-palmitoyl-2-oleoyl-phosphatidylethanolamine, 30% 1-palmitoyl-2-oleoyl-phosphatidylglycerol (Avanti Polar Lipids #850757 & #840457), 5 mg/mL in n-nonane (Alfa Aesar #A16177-AE) to form bilayers. Single channels were inserted by addition of protein reconstituted liposomes to the ‘cis’ side of the bilayer, and current was recorded at -200 mV holding potential (“cis” side defined as zero voltage) in symmetrical solutions containing 300 mM NaF, 10 mM NMG-Cl (Sigma-Aldrich #66930), 10 mM HEPES pH 7.0. Recordings were low-pass filtered at 160 Hz, digitized at 1.25 kHz, and analyzed in AXON Clampfit 10 (Molecular Devices Inc.). Channel conductance was measured from the difference between open vs blocked current in each recording. All recordings shown in figures are representative of behavior seen on many channels in multiple reconstitutions.

To calculate the dwell time distributions of open and closed channels, we consider a bimolecular binding scheme with rate constants of Na^+^ binding association *k*_*on*_ and dissociation *k*_*off*_:

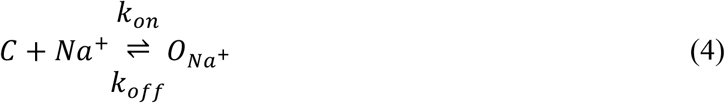

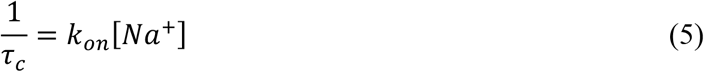

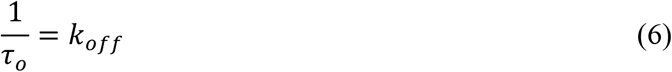

Where *O*_*Na+*_ represents the Na^+^ bound open channel.

### Subunit exchange assay via bulk FRET

The protein is labeled with either a donor-(Cy3-maleimide) or acceptor-(Cy5-maleimide) fluorophore or co-labeled with a mixture of both (in a 1:5 ratio of Cy3 to Cy5) and reconstituted into EPL membranes at 0.5 μg/mg. Donor- and acceptor labeled proteoliposomes are then mixed matching the measured co-labeled ratio and fused together using repeated freezing and thawing (F/T) cycles. FRET is measured in a fluorometer (FL3-22, Horiba) by exciting Cy3 at 534 nm, measuring the emission intensity, *I*_*Cy3*_, at 552 nm and *I*_*Cy5*_, at 655 nm and calculating the ratiometric FRET as follows:

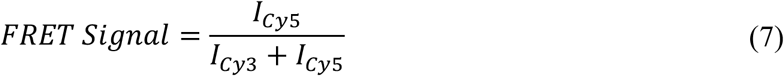

### Lipid mixing

Lissamine RhodamineB (RhB, acceptor) and 7-nitro-2-1,3-benzoxadiazol-4-yl-(NBD, donor) labeled PE lipids (Avanti Polar Lipids #810150 & #810145) were spiked in while preparing EPL liposomes at 1:40 for NBD-PE and 1:8 for RhB-PE. Donor- and acceptor labeled liposomes are then mixed in a 1:5 ratio and fused together using repeated freezing and thawing (F/T) cycles. FRET is measured in a fluorometer (Horiba) by exciting NBD at 465 nm, measuring the emission at *I*_*NBD*_ at 552 nm and *I*_*RhB*_ at 655 nm and calculating the ratio metric FRET via Equation 7.

### Single-molecule TIRF microscopy

Photobleaching experiments were carried out as described previously (*8*). Imaging was performed on an objective based total internal reflection fluorescence (TIRF) microscope, set-up for the imaging of single molecules. After loading the custom build slide onto the microscope, 30 μL of sample was loaded onto the lane and the flow cell was washed multiple times with dialysis buffer to remove any excess proteoliposomes not bound to the glass. The number of spots corresponding to proteoliposomes with labeled proteins was kept at a low density, less than 300 spots in the imaging field, by diluting the samples with dialysis buffer if needed. The total number of spots imaged for each measurement was at least 300 spots. Before imaging, the sample was passed through a 0.4 μm polycarbonate membrane (Whatman) using a LiposoFast-Basic (Avestin) extruder.

### Photobleaching data analysis and dimerization *K*_*D*_ estimation

The analysis of the photobleaching traces was carried out as described previously. Image files were analyzed using a MATLAB-based CoSMoS analysis program (*34*). Fluorescent spots were automatically selected by the image analysis software on the criteria of intensity and further selected via criteria (i.e. spots that were overlapping or at the edge of the field were excluded). Selected spots selected were analyzed using a 4 × 4 pixel area of interest (AOI) centered around the peak. The total pixel intensity within each AOI was integrated as a function of time, and then the intensity traces were examined for step-like decreases in intensity indicating irreversible photobleaching of fluorophores. The probabilities of single (*P*_*1*_), double (*P*_*2*_), and three or more steps (*P*_*3+*_) were calculated. For fitting, we followed a fitting protocol that has been developed by the lab and is described in detail elsewhere (*35*). The experimentally obtained photobleaching distributions were fitted to modeled photobleaching distributions using least-squared analysis by performing iterative fitting of *P*_*1*_, *P*_*2*_, *P*_*3+*_ vs. *χ*_*reconstituted*_ photobleaching data while varying *P*_*fluor*_ and *P*_*bg*_ for estimating the labeling yield via the WT dimer or varying *K*_*D*_ for stability estimations. The resulting *Normalized SSR*^*-1*^ is a probability distribution over the parameter space, where areas of higher probability are in better agreement with the experimental data than areas of lower probability. Using this distribution, we were able to infer an optimal point estimate as well as to quantify the uncertainty of all model parameters (*35*).

### Modeling and fitting cation linkage to dimerization stability

The analysis of cation linkage was carried out via Scientist® 2.0 from Micromath®. From the obtained *K*_*D*_ of the best fit, we calculated the fraction of dimer *F*_*Dimer*_ using

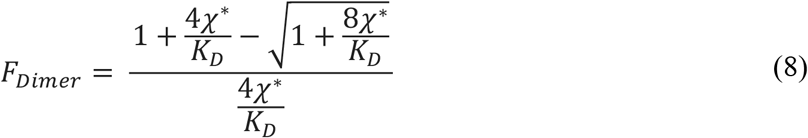

and *χ* = 1 × 10^−9^ to 1 × 10^−5^ subunit/lipid. The data was then fit to the following binding model using least-squared fitting:

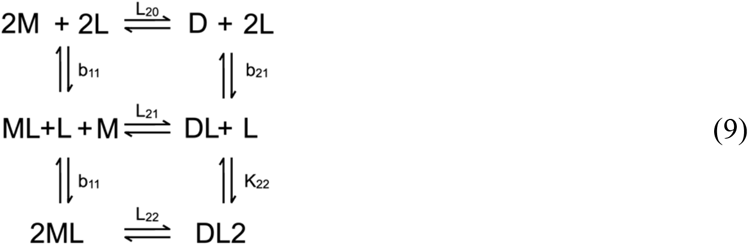

Using the following input:

~~~
// MicroMath Scientist Model File
IndVars: L,Mtot
DepVars: FDimer
Params: b11,b21,L20,K22
b22=b21*K22
M=(-
(1+b11*L)+((1+b11*L)^2+8*L20*(1+b21*L+b22*L^2)*Mtot)^0.5)/(4*L20*
(1+b21*L+b22*L^2))
M2=L20*M^2
M2L=L20*b21*M^2*L
M2L2=L20*b22*M^2*L^2
Fdimer=(2*M2+2*M2L+2*M2L2)/Mtot
0<M<Mtot
~~~

Where *Mtot* = *χ* and *L* is the Na^+^ concentration. To test sub-reactions of this system, parameters such as K22 or b11 were constrained to 0. To model sub-reactions of the system, we now used the simulation tool of Scientist® and the same model.

### MD simulations of Fluc in the monomer, dimer and dissociated dimer states

The simulations reported in this study used two lipid bilayers of identical composition but different dimensions, prepared using insane.py (*36*) based on the coarse-grained (CG) representation in the MARTINI 2.2 forcefield (*37*). Both bilayers are mixtures of POPE and POPG lipids, in a 3:1 ratio, initially immersed in a 150 mM NaCl solution. The smaller system is approximately 10.0 × 10.0 × 8.7 nm in size, and comprises 336 lipids, 3378 water particles, and 118 and 34 Na^+^ and Cl^-^ ions, respectively; the larger system is approximately 20.2 × 20.2 × 9.6 nm, and comprises 676 lipids, 16380 water particles, and 536 and 198 Na^+^ and Cl^-^ ions, respectively. To equilibrate each of these systems, we carried out an MD simulation of 50 μs at 1 atm and 303 K, using GROMACS 2018.8 (*38*). After verifying equilibration (system size, lipid mixing, etc.), the smaller bilayer was used to examine the membrane morphology associated with monomeric and dimeric Fluc; the larger bilayer was used to simulate the self-assembly of dissociated dimers.

CG models of monomeric and dimeric Fluc were based on Protein Data Bank entry 5NKQ and were constructed with a customized version of martinize.py (*36*). Elastic networks to sustain the secondary and tertiary structures of the protein were created using lower and upper cut-off distances of 0.5 and 0.9 nm, respectively, a force constant of 500 kJ mol^-1^ nm^-2^, and default values for the elastic-bond decay factor and decay power (-es 0 -ep 0). Glu88 was set as protonated due to its close proximity to a F^-^ ion in the experimental structure; all other ionizable residues were set at their default protonation state at pH 7. To simulate the monomer and dimer, the protein structures were embedded into the smaller POPE/POPG lipid bilayer, as captured in the last snapshot of the 50-μs trajectory used for equilibration. To do so, the protein structures were superposed on the bilayer at its center, and lipid, solvent, and ions within 3 nm from any protein particle were removed; the number of Na^+^ and Cl^-^ ions was then adjusted to increase the ionic strength to 300 mM while preserving a total net charge of zero. Following a series of short MD simulations to equilibrate the protein-lipid interface, a trajectory of 50 μs was calculated for each system, analogous to that calculated for the pure POPE/POPG lipid bilayer. The data reported in **Figures 5-6** derives from these trajectories, analyzed using the MOSAICS suite (*39*).

The dissociated dimer was simulated in the larger bilayer. To create this simulation system, a monomer was first embedded in the center of the membrane using the same protocol described above for the smaller bilayer. Following equilibration, a second monomer was inserted in the same membrane, in four different locations and orientations, at least 5 nm away from the monomer at the center. After equilibration, each of the four systems was simulated for 30 μs. To minimize the influence of direct monomer-monomer interactions in the outcome of these simulations, while simultaneously limiting the extent to which the two monomers can drift apart, the standard energy function was supplemented with an additional potential energy term acting on the minimum distance between the two monomers, or *d*_min_; to calculate this distance during runtime, all backbone CG particles in the transmembrane region of each protein were considered (30 per monomer). Specifically, this potential energy term is *E’* = 0.5 × *K* × (*d* - *d*_*min*_)^2^ if *d*_*min*_ ≤ 1.3 nm or *d*_*min*_ ≥ 5.5 nm with *K* = 500 kJ mol^-1^ nm^-2^. Note that for any other value of *d*_*min*_ this additional term is inactive i.e. *E’* = 0. Calculations of *d*_*min*_, *E’* and the associated atomic forces were carried out using PLUMED 2.2.5 (*40*).

### Statistical Analysis

Statistical analysis in each required experiment was performed using either unpaired Student’s *t* test (two-tailed distribution) or analysis of variance (ANOVA) test with GraphPad Prism version 9 (one-way ANOVA and Uncorrected Fisher’s LSD). Details of statistical analyses, including sample sizes and biological replications, are provided in the figure legends.

## Acknowledgments

The authors thank Dr. Benjamin McIlwain for preliminary electrophysiology experiments for N43S Fluc., Dr. Nathan Bernhardt for his assistance with data analysis, Dr. Rahul Chadda for his guidance on the FRET studies and the Robertson lab members for useful discussion.

## Funding

National Institutes of General Medical Sciences, National Institutes of Health grant R01GM120260 (ME, JLR)

National Institutes of General Medical Sciences, National Institutes of Health grant R35GM128768 (RBS)

Millipore Sigma Predoctoral Award (ME)

American Heart Association grant 23PRE1013841 (ME)

Intramural Research Program of the National Heart, Lung and Blood Institute (NHLBI),

National Institutes of Health (NIH) (EO, JDFG)

NIH High-Performance Computing System Biowulf (EO, JDFG)

## Author contributions

Conceptualization: ME, JDFG, JLR

Methodology: ME, EAO, RBS, JDFG, JLR

Investigation: ME, EAO

Visualization: ME, JDFG, JLR

Supervision: RBS, JDFG, JLR

Writing—original draft: ME, JDFG, JLR

Writing—review & editing: ME, JDFG, JLR

## Data and materials availability

All data are available in the main text or the supplementary materials.

