## Supplementary Figures & Table for "Dimerization mechanism of an inverted-topology ion channel in membranes"

**This PDF file includes:**

Figs. S1 to S7
Table S1

**Other Supplementary Materials for this manuscript include the following:**

Data S1

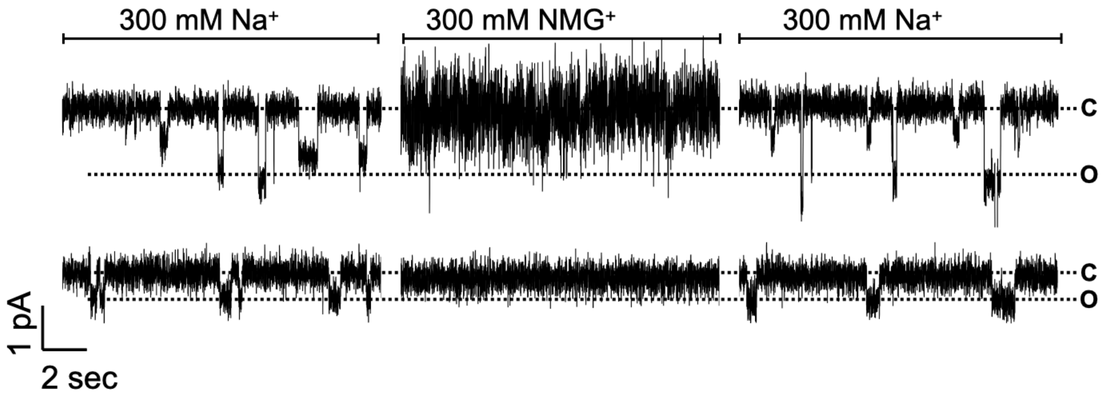


**Fig. S1. Na$^+$ dependency is independent of subpopulation.** Na$^+$ exchange in single channel recording. Representative traces of single channel recordings of active N43S-Cy5 reconstituted at 0.05 µg/mg in EPL, inserted into a 2:1 POPE/POPG bilayer and recorded at -200 mV. The upper lines mark the closed and the lower lines mark the open state of the channel. Single channel was recorded in 300 mM NaF buffer in cis and trans chamber. Then buffer was switched to 300 mM NMGF in cis chamber. Buffer was then switched back to 300 mM NaF in cis chamber. Upper panel shows a channel with high current amplitude, lower panel shows a channel with lower current amplitude.

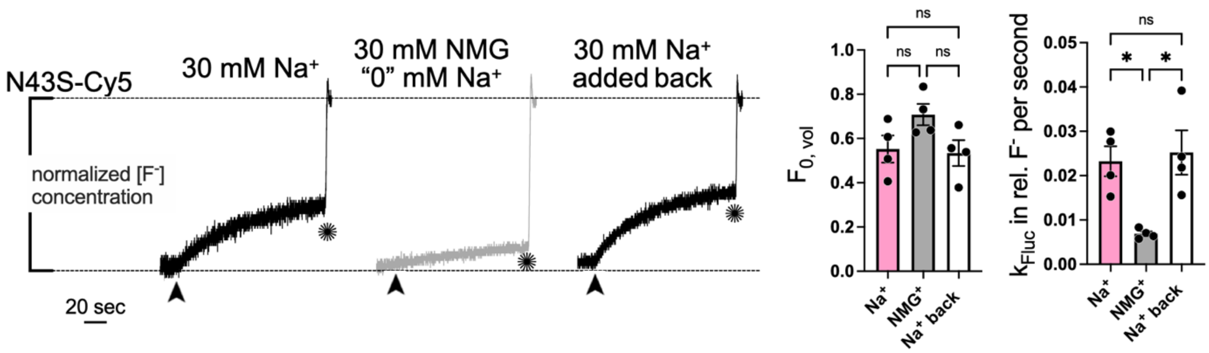


**Fig. S2. N43S-Cy5 shows $Na^+$ dependent fluoride transport in macroscopic transport assay.** Representative traces of normalized fluoride concentration of external solution over time with 30 mM $Na^+$ present (black), replaced by 30 mM $NMG^+$ via dialysis (grey, henceforth called "0" mM $Na^+$) and 30 mM $Na^+$ added back (grey). Right: Raw traces are fitted with $Y = (1-F_0)*(1-\exp(-(k_{Fluc}+k_{leak})*x))+F_0*(1-\exp(-k_{leak}*x))$, where $k_{leak} = 8.5 \times 10^{-5}$ rel. $F^-$ per second. $F_0$ and $k_{Fluc}$ for all three samples reported as mean ± SE, n = 3. The fractional volume of inactive vesicles $F_0$ of all three samples are similar ($Na^+$ vs. $NMG^+$: $p = 0.1783$, $Na^+$ vs. $Na^+$ back: $p = 0.9710$, $NMG^+$ vs. $Na^+$ back: $p = 0.1265$, Ordinary one-way ANOVA), while the rates of fluoride transport $k_{Fluc}$ of samples with $Na^+$ are significantly different than the one without ($Na^+$ vs. $NMG^+$: 0.0225, $NMG^+$ vs. $Na^+$ back: $p = 0.0122$, ordinary one-way ANOVA) but there is no significant change in transport rate of $Na^+$ control vs. $Na^+$ added back ($p = 0.9167$, ordinary one-way ANOVA).

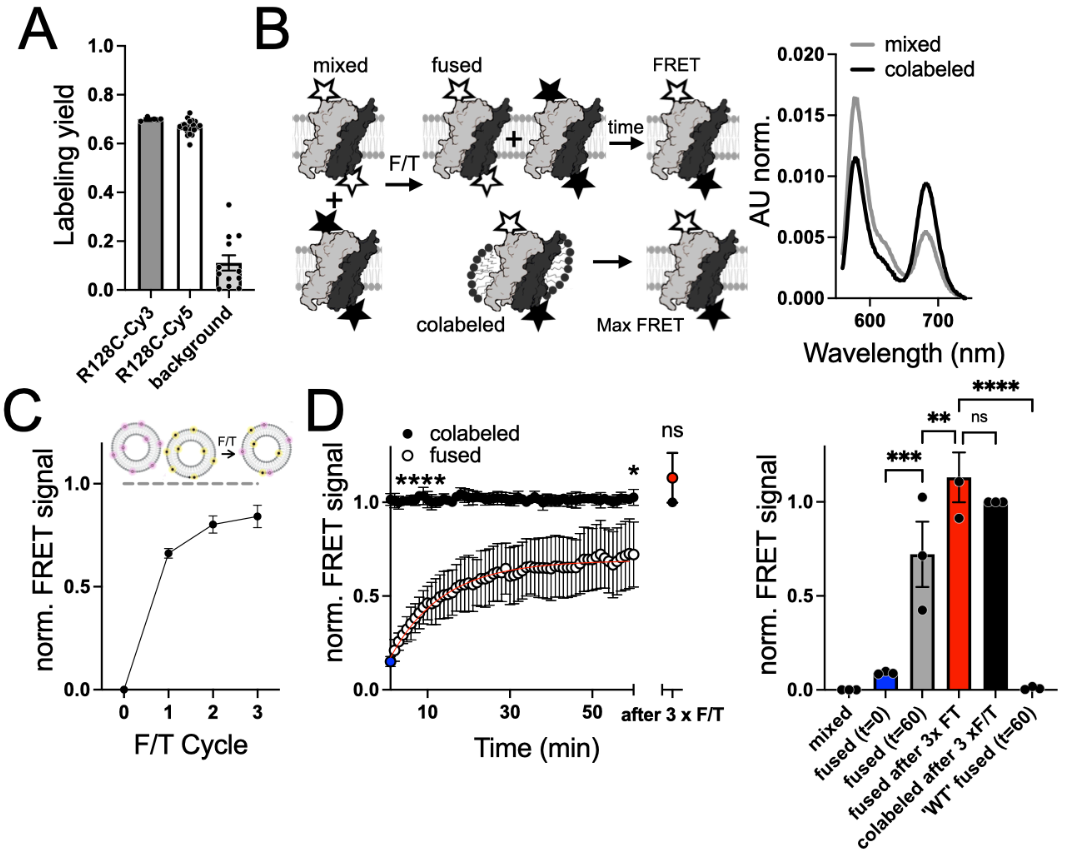


**Fig. S3. N43S shows rapid subunit exchange.** (**A**) Labeling yields of N43S labelled at R128C for Cy3 and Cy5 as well as background. (**B**) Cartoon of bulk FRET subunit exchange assay. Donor- and acceptor labeled Fluc (N43S-Cy3 and N43S-Cy5, respectively) are reconstituted separately, and liposomes are mixed (mixed) and fused using repeated F/T cycles. Upon fusion, subunits can freely exchange leading to an increase in FRET signal over time that approaches a maximal FRET signal determined by colabeling the protein with both fluorophores before reconstitution (co-labeled). Right side: Representative raw traces of fluorescence intensity over wavelength for mixed and colabeled. (**C**) Fusion of liposomes is complete after 2 x F/T cycles. Normalized FRET signal from fusing NBD- or RhB-labeled liposomes as a function of F/T cycles. Normalized to colabeled control where both, donor and acceptor are incorporated into liposomes ('positive control', dashed line). Data are reported as mean ± SE, n = 2. (**D**) FRET signal increase of N43S-Cy3 fused with N43S-Cy5 shows rapid subunit mixing in in the absence of $Na^+$ ("0" mM $Na^+$). 'Co-labeled' represents the maximal FRET. 'Mixed' is the FRET signal after mixing but before fusion of the liposomes. 'Fused' shows the FRET signal after liposome fusion via 1 x F/T and incubation at RT for one hour. The data is fit with a single exponential (red line). After 3 x F/T shows the FRET value after performing two additional F/T cycles of the sample. Data are reported as mean ± SE, n = 3. P values: fused ($t$ = 0, blue) vs. colabeled ($t$ = 0): <0.0001, fused ($t$ = 60) vs. colabeled ($t$ = 60): 0.0357, fused after 3 x F/T vs. colabeled after 3 x F/T: 0.3246. Right side: The fused sample after 1 x F/T and 60 min incubation (fused $t$ = 60) shows significantly higher FRET compared to the fused sample after 1 x F/T but before incubation (fused $t$ = 0) and lower FRET signal compared to after 3 x F/T but no difference compared to the co-labeled control (colabeled after 3 x F/T). 'WT'-R128C does not show subunit exchange as the FRET signal does not increase after 60 min incubation after fusion (‘WT’ fused $t$ = 60). Data are reported as mean ± SE, n = 3 ($p$-values: fused ($t$ = 0) vs. fused ($t$ = 60): 0.0003, fused ($t$ = 60) vs. fused after 3 x F/T: 0.0073, fused after 3 x F/T vs. colabeled after 3 x F/T 0.3246, fused after 3 x F/T vs. ‘WT’ fused ($t$ = 60): <0.0001, Ordinary one-way ANOVA).

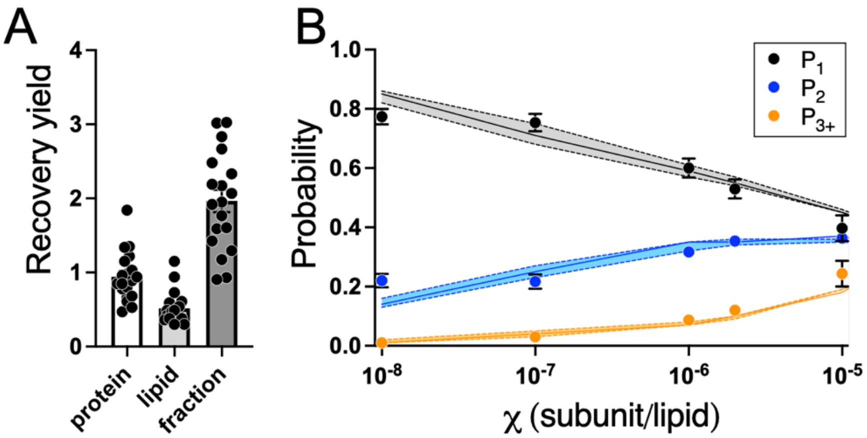


**Fig. S4. Fitting of N43S-Cy5 photobleaching data.** (**A**) Protein to lipid mole fraction recovery yield after reconstitution. Protein and lipid recovery individually as well as the fraction of both. (**B**) N43S-Cy5 photobleaching data were fit using the estimated values $P_{fluor}$ = 0.67 and $P_{bg}$ = 0.11 to estimate macroscopic $K_D$ values for each ion concentration. Shown are the experimental $P_1$, $P_2$, $P_{3+}$ distributions as well as the best fit mean ± SE (shaded area).

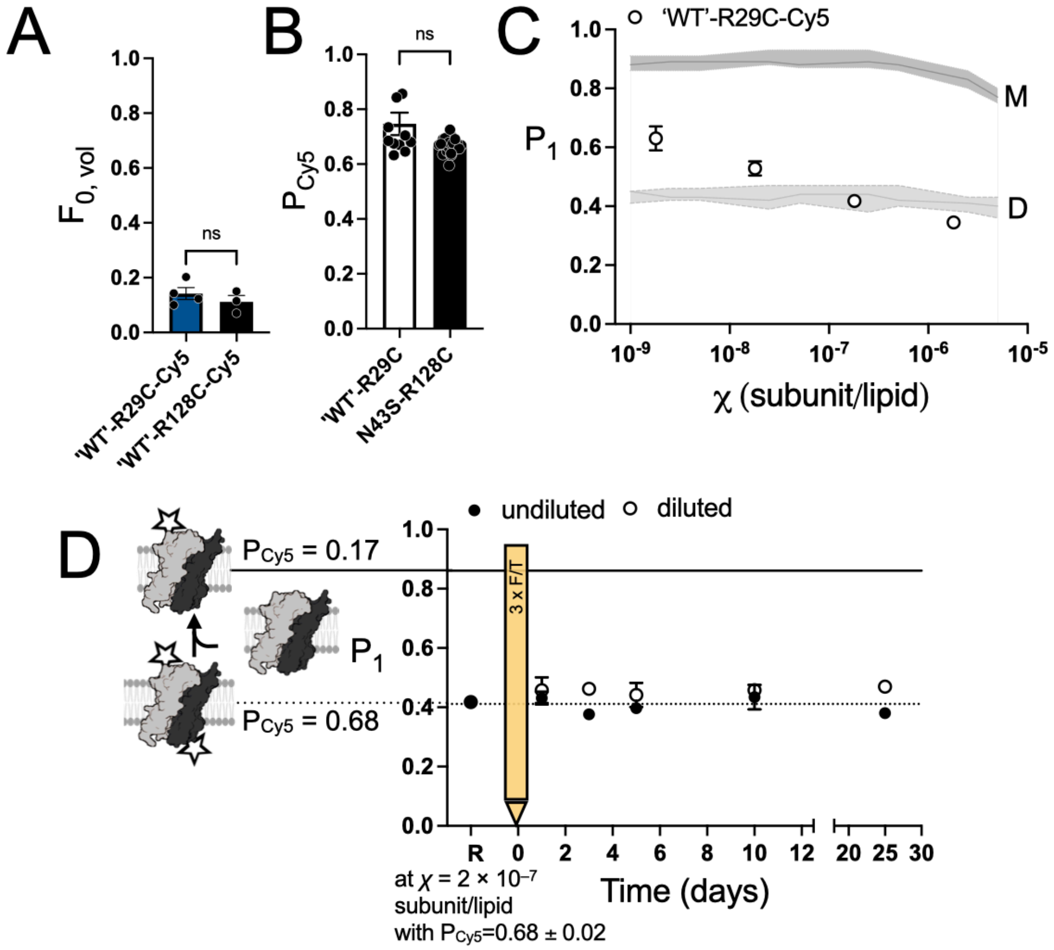


**Fig. S5. Fluc WT is a kinetically trapped dimer.** (**A**) 'WT'-R29C site specifically labelled with Cy5 shows comparable labeling yield to N43S-R128C-Cy5. (**B**) 'WT'-R29C-Cy5 is functional with transport rates that exceed the response time of the electrode and $F_0$ values similar to previously reported values as well as when labelled at R128C. (**C**) WT'-R29C-Cy5 is dimeric over $\chi = 2 \times 10^{-9}$ subunit/lipid to $\chi = 2 \times 10^{-6}$ subunit/lipid. Shown are the photobleaching probabilities of a single step ($P_1$) vs $\chi$ for 'WT'-R29C-Cy5 (open circles), and theoretical monomer (M) and dimer (D) controls modeled using the experimental labeling yields $P_{Cy5}$ and $P_{bg}$ following Chadda et al., 2016. Data are reported as mean ± SE n = 3. (**D**) Dilution study of reconstituted labeled protein with unlabeled protein. Post-freeze/thaw (F/T) time course of 'WT'-R29C-Cy5 at mole fraction $\chi = 2 \times 10^{-7}$ subunit/lipid diluted 1/4 with unlabeled protein (open circles) and undiluted control sample (closed circles) shows no change. "R" represents the original high-density reconstituted sample at $\chi = 2 \times 10^{-7}$ subunit/lipid, prior to dilution. The F/T process is indicated by the yellow bar. The dotted and solid line report on the modeled $P_1$ for $P_{Cy5} = 0.68$ and $P_{Cy5} = 0.17$, respectively. Data reported as mean ± SE, n = 2-3.

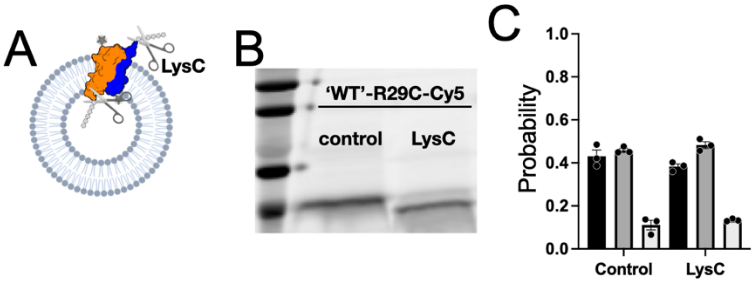


**Fig. S6. His-Tag removal does not influence photobleaching distribution.** (**A**) Cartoon representation of His-Tag cutting via LysC. Of notion is the antiparallel topology of Fluc that necessitates repeated F/T and extrusion cycles to make the protein subunits accessible to LysC which cannot permeate the membrane. (**B**) SDS-PAGE image of control and protein cut with LysC. The LysC sample shows a shift in the band indicative of his-tag cleavage. (**C**) Photobleaching distributions of 'WT'-R29C-Cy5 control and LysC-treated sample at $\chi = 2 \times 10^{-7}$ subunit/lipid show no significant difference ($p = 0.2432$, $\chi^2$ test).

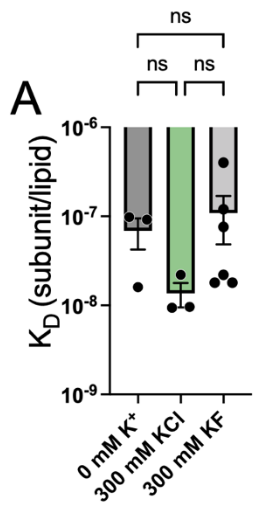


**Fig. S7. $K^+$ and anions do not significantly influence dimer stability.** Fitted $K_D$ following (*35*) for 0 mM $K^+$, 300 mM KCl on 30 mM $Na^+$ background (300 mM KCl), 300 mM KF on 30 mM $Na^+$ background (300 mM KF) shows no significant difference between all samples (*p*-values: 0 mM $K^+$ vs. 300 mM KCl: 0.7579, 0 mM $K^+$ vs. 300 mM KF: 0.13, 300 mM KCl vs. 300 mM KF: 0.0725, Ordinary one-way ANOVA).

| Model | Parameter | Estimated Value | Standard Deviation | 95% Range (Univar) | |
|---|---|---|---|---|---|
| $Na^+$ binding to single site at dimer (Model 1) | b21 | 60.42 | 9.65 | 41.46 | 79.38 |
| | L20 | 2.81 x $10^7$ | 0.28 x $10^7$ | 2.27 x $10^7$ | 3.35 x $10^7$ |
| $Na^+$ binding to monomer and dimer (Model 2) | b11 | -0.46 | 0.28 | -1.01 | 0.09 |
| | b21 | 46.52 | 11.21 | 24.49 | 68.55 |
| | L20 | 3.03 x $10^7$ | 0.32 x $10^7$ | 2.41 x $10^7$ | 3.66 x $10^7$ |
| 2 $Na^+$ binding to dimer (Model 3) | b21 | 47.02 | 11.11 | 25.18 | 68.85 |
| | L20 | 3.04 x $10^7$ | 0.32 x $10^7$ | 2.41 x $10^7$ | 3.66 x $10^7$ |
| | K22 | 1.19 | 0.92 | -0.62 | 3.00 |
| $Li^+$ binding to single site at dimer | b21 | 72.24 | 8.40 | 55.71 | 88.77 |
| | L20 | 2.97 x $10^7$ | 0.21 x $10^7$ | 2.56 x $10^7$ | 3.38 x $10^7$ |
| $K^+$ binding to single site at dimer | b21 | 2.36 | 0.28 | 1.80 | 2.91 |
| | L20 | 1.58 x $10^8$ | 0.05 x $10^8$ | 1.48 x $10^8$ | 1.68 x $10^8$ |

**Table S1. Fitting parameters of cation binding models.**

**Data S1. Source data file for Fig. 1 to Fig. 4.**
